# Comparative toxicity of menthol- and tobacco-flavored electronic cigarette constituents causing inflammation, epithelial barrier dysfunction, and nicotinic acetylcholine receptor modulation in the absence of nicotine

**DOI:** 10.1101/2025.06.30.662345

**Authors:** Vidhi S. Pandya, Arni Bhatnagar, Kirby J. Beck, Thivanka Muthumalage

**Affiliations:** School of Health Sciences, Purdue University, West Lafayette, IN, USA

**Author notes:** **Corresponding Author:** Thivanka Muthumalage, PhD, Address: Purdue University, Lilly Hall of Life Sciences, 915 Mitch Daniels Blvd, LILY 2-121, West Lafayette, IN 47907, Lab.

**Keywords:** Inflammation, electronic cigarettes, flavoring, electronic nicotine delivery systems, tobacco, menthol, acute lung injury, chronic obstructive pulmonary disease

## Abstract

**Background:** Menthol and tobacco-flavored nicotine delivery systems (ENDS) are widely used as safer alternatives to combustible cigarettes. These flavored products include constituents such as propylene glycol/vegetable glycerin (PG/VG), benzoic acid, acetoin, L-menthone, 98% menthone, 2-isopropyl-N,2,3-trimethylbutanamide (WS-23), vanillin, and carvone. However, little is known about the potential adverse effects of the constituents in these flavored products.

**Rationale and hypothesis:** We hypothesized that exposure to common constituents in tobacco- and menthol-flavored ENDS constituents could elicit a lung-injurious response mediated by nicotinic acetylcholine receptor (α-nAChR or CHRNA) modulation.

**Methods:** Human bronchial epithelial cells, BEAS-2B, cells were treated with commonly used menthol and tobacco constituents on trans well inserts. Transepithelial barrier resistance (TEER) and millivolts (mV) across epithelial cells were measured over a 24-hour time. To assess the elicited inflammatory response, cytokines IL8 and IL6 were quantified in the conditioned media. Cytotoxicity caused by these constituents was evaluated by acridine orange/propidium iodide (AO/PI) staining of the cells after 24 hrs. alpha nicotinic receptor protein abundance (α1, α4, α5, and α7) was quantified by immunoblotting.

**Results:** Epithelial integrity was decreased over time with a significant decrease in TEER and voltage by ENDS constituents. A significant increase in IL6 in conditioned media was observed in PG/VG, carvone, and WS-23 treated cells. Carvone-treated cells also elicited significantly elevated IL8 in conditioned media. Further, increased α1, α4, α5, and α7 nAChR were seen in cells treated with PG/VG, Acetoin, Carvone, and WS-23.

**Conclusion:** These findings suggested that common constituents in menthol- and tobacco-flavored ENDS induce lung inflammation, epithelial barrier dysfunction, and lung injury. Further, our data implicate potential lung disease pathogenesis via nAChR modulation-mediated inflammation by exposure to these ENDS constituents, even in the absence of nicotine.

## Background

The usage of electronic nicotinic delivery systems (ENDS) or electronic cigarettes has increased in the past few decades. ENDS are often used as a smoking cessation product or a safer alternative to cigarettes. However, these ENDS products have also been used by populations who had never used tobacco products. Electronic Nicotine Delivery Systems (ENDS) consumption has increased by 22.1% from 2011-2020 in youth and by 2.6% from 2012-2019 (1). The popularity of menthol and tobacco flavors is more prevalent among adults, around 22.3-31.8% (1). Menthol flavor use has increased among youth participating in the PATH study from Wave 2 to Wave 5 (2).

Additionally, the flavor ban by the FDA in 2020 on flavors except menthol and tobacco in closed pods had also been a promoter of the increased use of these flavoring agents (3) **(Table 1)**. These were exclusive for changeable pod devices with E-liquids, not for any disposable ENDS products. The use of menthol is more popular as it provides a soothing sensation upon inhalation and has antitussive properties (4). Along with those, higher concentrations of menthol are also associated with analgesic effects, which could lead to an increased consumption of E-cigarettes and nicotine (4, 5).

**Table 1:**
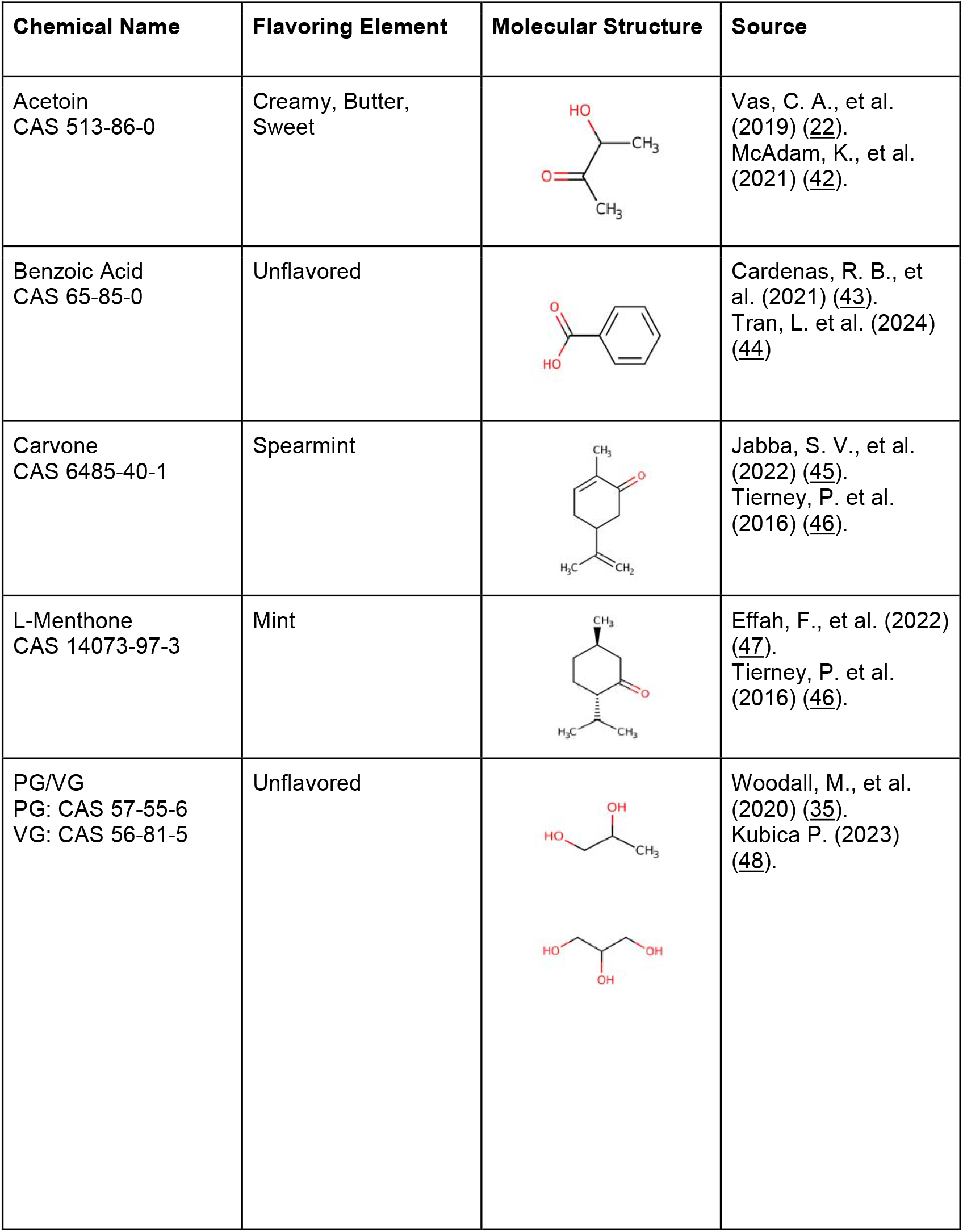
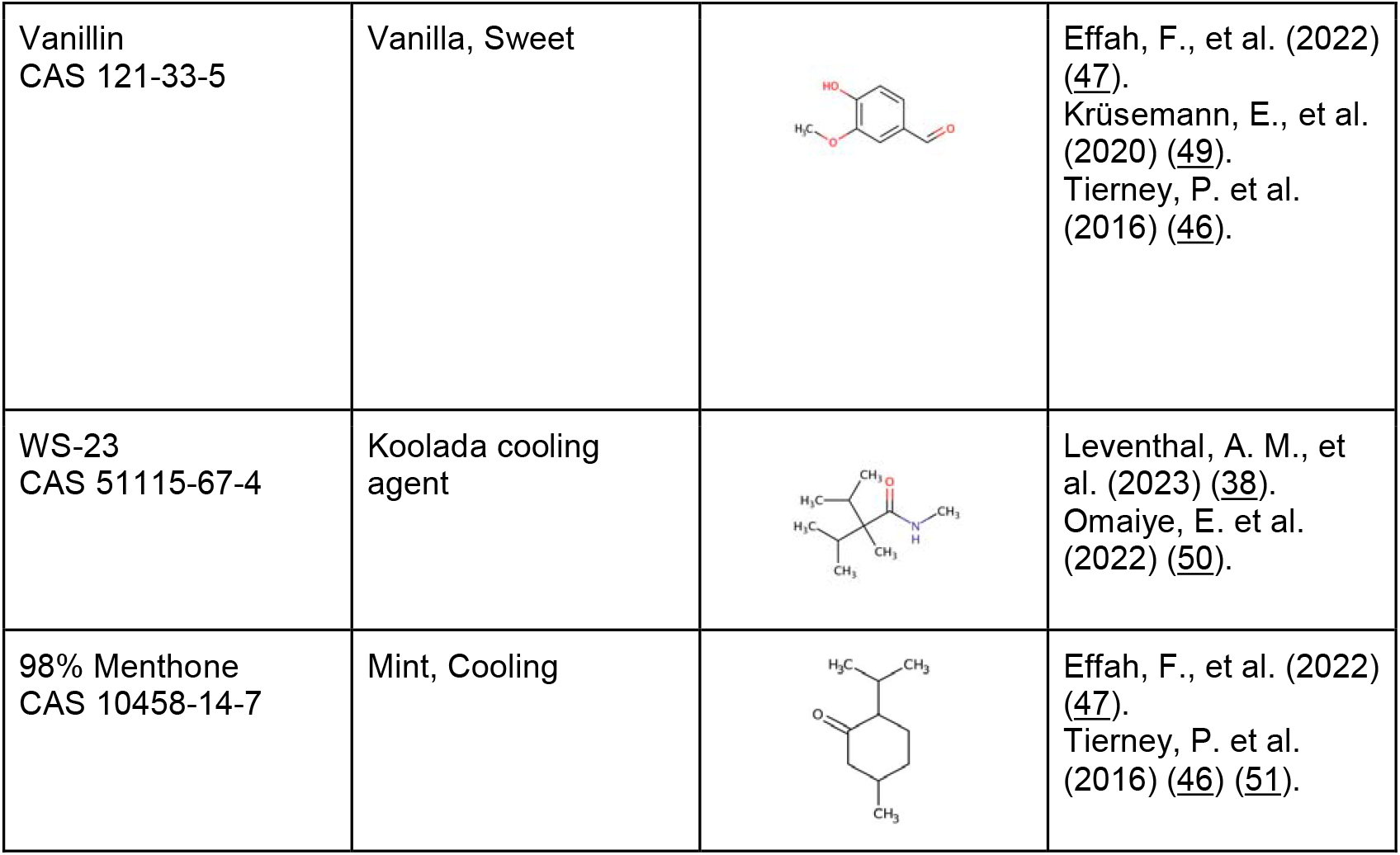
Constituents of menthol- and tobacco-flavored ENDS evaluated in this study.

| Chemical Name | Flavoring Element | Molecular Structure | Source |
| --- | --- | --- | --- |
| Acetoin<br>CAS 513-86-0 | Creamy, Butter,<br>Sweet |  | Vas, C. A., et al. (2019) (22).<br>McAdam, K., et al. (2021) (42). |
| Benzoic Acid<br>CAS 65-85-0 | Unflavored |  | Cardenas, R. B., et al. (2021) (43).<br>Tran, L. et al. (2024) (44) |
| Carvone<br>CAS 6485-40-1 | Spearmint |  | Jabba, S. V., et al. (2022) (45).<br>Tierney, P. et al. (2016) (46). |
| L-Menthone<br>CAS 14073-97-3 | Mint |  | Effah, F., et al. (2022) (47).<br>Tierney, P. et al. (2016) (46). |
| PG/VG<br>PG: CAS 57-55-6<br>VG: CAS 56-81-5 | Unflavored |  | Woodall, M., et al. (2020) (35).<br>Kubica P. (2023) (48). |
| Vanillin<br>CAS 121-33-5 | Vanilla, Sweet |  | Effah, F., et al. (2022) (47).<br>Krüsemann, E., et al. (2020) (49).<br>Tierney, P. et al. (2016) (46). |
| WS-23<br>CAS 51115-67-4 | Koolada cooling agent |  | Leventhal, A. M., et al. (2023) (38).<br>Omaiye, E. et al. (2022) (50). |
| 98% Menthone<br>CAS 10458-14-7 | Mint, Cooling |  | Effah, F., et al. (2022) (47).<br>Tierney, P. et al. (2016) (46) (51). |

Tobacco-induced diseases such as chronic obstructive pulmonary disease (COPD), asthma, and idiopathic pulmonary fibrosis (IPD) are mediated via nicotinic receptors (nAchRs or CHRNAs) (6). Nicotine in tobacco interacts with *α*7 nAchR, mediating lung injury(7). Similarly, nicotine from electronic cigarettes is also known to bind to cholinergic nicotinic receptors, mediating addictive properties by activating the mesolimbic reward system(8). However, it is unknown if other constituents in ENDS can modulate nicotinic receptors. To investigate the knowledge gap, we hypothesized that non-nicotine constituents in ENDS can modulate CHRNAs that mediate lung pathogenesis. Thus, we identified the key flavoring chemicals in Table 1 and assessed their ability to induce lung injury via CHRNAs. These receptors play a significant role in pulmonary epithelial cells, especially when exposed to foreign chemicals (6). Nicotine from E-liquids is known to interact with many nAChRs and has been implicated in regulating the immune system, leading to inflammation (6). These receptors also possess metabotropic functions in immune cells. CHRNAS *a*7, *a*5, *a*4, and *a*1 were analyzed due to their prominent roles in the bronchial epithelial airway (9). nAChR *a7-dependent* pathways also played a role in the prognosis for asthma and COPD (9). Additionally, the activation of *a*5 is linked with cell proliferation through the TRPC3 channels (9). Some of these nAChRs have also displayed metabotropic signaling properties when bound to GTP-binding proteins that regulate cytokine expression (10).

Interleukin-6 (IL6) is a known regulator of lung inflammation in addition to other inflammatory cytokines such as TNF*α*, IL1ß, and GM-CSF (11). These cytokines recruit other leukocytes, inducing oxidative stress and lung inflammation (12) (13). Increased cytokine levels have been observed in asthmatic patients’ bronchoalveolar lavage fluid (BALF) compared to healthy non-smoking patients and stable non-asthmatic and asthmatic patients on mechanical ventilation (14). The induction of IL-8, often synthesized by TNF*α*, has also been linked to conditions such as COPD, acute respiratory distress syndrome (ARDS) (13), and asthma (15-17). Free IL-8 in bronchial tissue is also linked with patients experiencing severe asthma (18). The pro-inflammatory effects of IL-8 have previously been linked to a stress or injury response (19). Increased levels of IL-8 in BALF of many acute lung injury (ALI) patients have also been linked with an increase in mortality (20). Like IL-6, increased levels of IL-8 in BALF have also been observed in several asthmatic patients (21, 22). The risk of developing chronic obstructive pulmonary disease (COPD) is related to chronic inflammation in the lungs (23) through cytokine release and oxidative stress, which can lead to damaged lung parenchyma (24). Additionally, chronic inflammation is associated with the expression of NF-κB transcription factor that plays a vital role in inflammation-induced carcinogenesis (24). It is also linked to the expression of pro-inflammatory cytokines such as IL-6 and IL-8 (16).

Exposure to vaping constituents can elicit an immune response and cause lung epithelial injury. Tight junctions–such as claudins and occludins–adheren junctions, and desmosomes play a role in maintaining permeability and polarity in adjoining cells. Barrier dysfunction in the epithelium has been linked with COPD, asthma, and pulmonary fibrosis (25). The presence of the tight junctions on the apical side of the trans wells regulates paracellular transport between adjacent cells and adherens which aids in cell-to-cell adhesion (25). The utilization of the air-liquid interface (ALI) in this study was best suited to mimic the respiratory epithelium and analyze barrier functionality in addition to the assessment of cytokine release (25).

To test our hypothesis, in this study, we assessed pulmonary toxicity of common flavoring chemicals and constituents in tobacco- and menthol-flavored ENDS–menthone, cooling agent (WS23), carvone, acetoin, vanillin, benzoic acid, and PG/VG–utilizing a bronchial epithelial ALI cell model. The study will provide insights into the comparative toxicity of flavoring constituents in ENDS and their engagement in modulating nicotinic receptors.

## Methods

### Scientific rigor and reproducibility

We applied a robust, unbiased experimental design, and data analysis approach throughout the study. We validated the methods and ensured reproducibility with repeated experiments. All methods are presented in detail with transparency. Results were reported and interpreted without bias. For all assays, laboratory-grade biological and chemical resources were purchased from commercial sources. Our methodologies, data, and results adhered to strict NIH reproducibility standards and scientific rigor.

### Cell Culture and treatments with flavoring chemicals

Human bronchial epithelial cells (BEAS-2B) (ATCC) were cultured on transwells (Corning #3402) in Gibco Dulbecco’s Modified Eagle Medium (DMEM) supplemented with 5% fetal bovine serum (FBS), 15 mM HEPES, 1% L-glutamine, and 1% antibiotic-antimycotic. Cells were incubated at 37º C and 5% CO_2_. Once cells reached 80% confluency, they were serum-deprived with 1% FBS, and treatments were performed at 90% confluency.

To perform flavoring chemical treatments, 100 mM stocks were made for L-Menthone (Thermo Scientific, Catalog: A13679.18), 98% Menthone (Thermo Scientific, Catalog: 125411000), (R)-(−)-Carvone (Thermo Scientific, Catalog: A13900.18), and Acetoin (Sigma Aldrich, Catalog: 40127-U) by diluting respected volumes of them in molecular grade Ethanol. For the solids such as Vanillin (Thermo Scientific, Catalog: A11169.22), Benzoic Acid (Fisher Scientific, Catalog: A65-500), and WS-23 (N-2,3-Trimethyl-2-isopropylbutanamide) (Sigma Aldrich, Catalog CDS003481), respected masses were diluted in molecular grade Ethanol to make 100 mM stocks. The working concentration of 100 µM was achieved through diluting the stocks in 1% FBS media. Equal parts of Propylene Glycol (Fisher Scientific, Catalog: P355-4) and Glycerol (Fisher Scientific, Cat G31-1) were used to make a 0.25% solution diluted in 1% FBS media. 100 µM of Ethanol and Hydrogen Peroxide (Fisher Scientific, Catalog: H325-500) were used as the vehicle controls to ensure that the amount used for the treatments was not cytotoxic. TNFD (10 ng/µL) (Gibco, PHC3015) was used as a positive control.

### Lung epithelial barrier integrity

Cells were grown on transwells until they reached a monolayer. Then, the EVOM^2^ Epithelial Volthommeter from World Precision Instruments was used to measure the transepithelial electrical resistance (TEER) and the voltage (mV) of the cells. Once the cells formed a monolayer with constant resistance, cells were serum-deprived overnight and treated with the chemicals of interest: L-menthone, 98% menthone, carvone, WS-23, vanillin, benzoic acid, PG/VG, and acetoin as described above. Resistance and voltage readings were obtained in 3 positions per transwell with the same treatment with STX2 electrode (chopsticks). An average was calculated for those three readings for each well and each treatment was performed in three wells. Voltage (part of the TEER calculation, V=IR) and resistance readings were obtained before any of the chemical treatments for all the plates, followed by readings at 6, 8, 20, and 24 hours after the treatments and also compared to untreated control. TEER was calculated by multiplying the surface area by the net resistance of the well.

### Cytotoxicity Assessment

Twenty-four hours post-treatment, the conditioned media from apical and basal membrane media were collected from each well for cytokine assessment. Cells were then trypsinized (0.25% Trypsin-EDTA, Gibco), and cell viability was assessed by AO/PI (Acridine Orange/Propidium Iodide) stain with live, dead, and total cell counts were obtained by DeNovix CellDrop automated cell counter (Denovix, DE, USA).

### Cytokine quantification by ELISA Assay

The collected conditioned media from apical layer media was used to analyze for the presence of Interleukin-6 (IL-6) (Invitrogen, Cat# KHC0061) and Interleukin-8 (IL-8) (Invitrogen, Cat# KHC0081) according to the manufacturer’s protocol by ELISA using Accuris Smart reader.

### Immunoblotting

Harvested cells from each treatment group were lysed with cOmplete protease inhibitor cocktail (Roche) and total protein was estimated by BCA protein assay (Thermofisher, Cat# 23225). An equal amount of protein (5-10 µg) were added to 10% SDS-PAGE gels along with a Precision plus protein standard (Biorad, Cat#1610376). Gel was transferred to PVDF membranes using Trans-Blot turbo (Biorad) system and the transfer was confirmed by Ponceau S stain (Thermofisher, A40000279). Blots were blocked with 5% non-fat dry milk in TBST (ASI, Cat# MB9696) for one hour at room temperature or overnight at 4C. The membranes were probed with specific primary antibodies: CHRNA1 (Abcam, Cat #AB308306), CHRNA4 (Abcam, Cat #AB124832), CHRNA5 (Abcam, Cat #AB259859), and CHRNA7 (Abcam, Cat #AB216485) overnight in a 4º C. The membranes were washed for 30 minutes with TBST and then incubated with HRP-conjugated goat anti-rabbit secondary antibody (Abcam, Cat #AB6721) for an hour at room temperature. After the membranes were washed 10 mins 3X, membranes were after chemiluminescence was detected by (SuperSignal West Femto Thermofisher, Cat# 34094) using the Bio-Rad ChemiDoc XRS+ system. Subsequently, the blots were gently stripped for 10 mins, washed, and re-probed with remaining primary antibodies and finally with loading control ß-actin (Abcam, Cat #AB8227). Densitometry analysis was done using Image Lab software (Biorad) to analyze band intensity. Data was normalized by loading control ß-actin.

### Statistical analyses

All statistical analysis tests were managed on GraphPad Prism (Version 10.3.1). P<0.05 was deemed statistically significant.

## Results

### Tobacco and menthol ENDS flavoring chemicals cause epithelial barrier dysfunction

All tested constituents–L-menthone, 98%menthone, carvone, WES-23, vanillin, benzoic acid, and PG/VG, caused a decrease in epithelial barrier integrity compared to pretreatment and untreated control **(Fig. 1)**. The resistance of the cells exposed to L-Menthone and 98% Menthone showed a decrease over time, but not statistically significant **(Fig. 1 A, B)**. In comparison, L-Menthone and 98% Menthone caused a more statistically significant reduction in voltage, with 98% Menthone showing more significance at the 6-hour time point when compared to L-Menthone **(Fig. 1 A, B)**. Carvone caused a significant decrease in voltage over time **(Fig. 1C)**. Similarly, cooling agent, WS23, also had a significant decline in mV over time. WS-23 caused a similar decrease in TEER but the voltage drop at 24-hour time point was statistically significant **(Fig. 1D)**. Acetoin and vanillin caused a significant decrease in both resistance and voltage over the course of 24 hrs with a slight improvement in mV during middle time points **(Fig. 1E, F)**. The primary vehicle component in e-liquids, PG/VG, caused a significant reduction in TEER and voltage at 24 hrs (**Fig. 1G**). Benzoic acid, used in nicotine salt products, caused a significant decrease in both resistance and voltage at 8, 20, and 24-hour time points **(Fig. 1H)**. Overall, either both TEER and mV or one of those parameters which is indicative of epithelial tight-junction health was adversely affected and declined over the 24-hour period compared to untreated controls.

**Figure 1:**
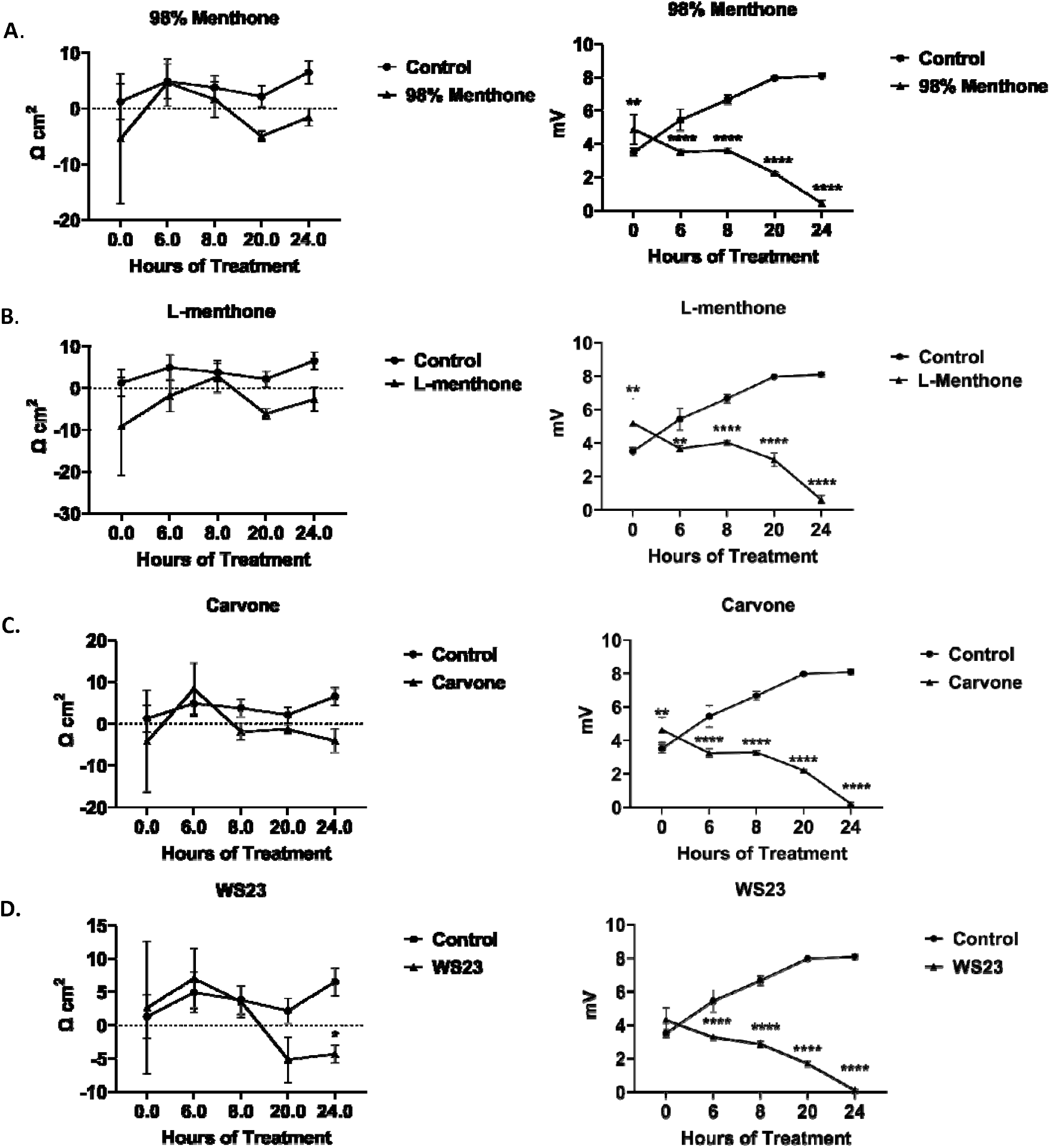
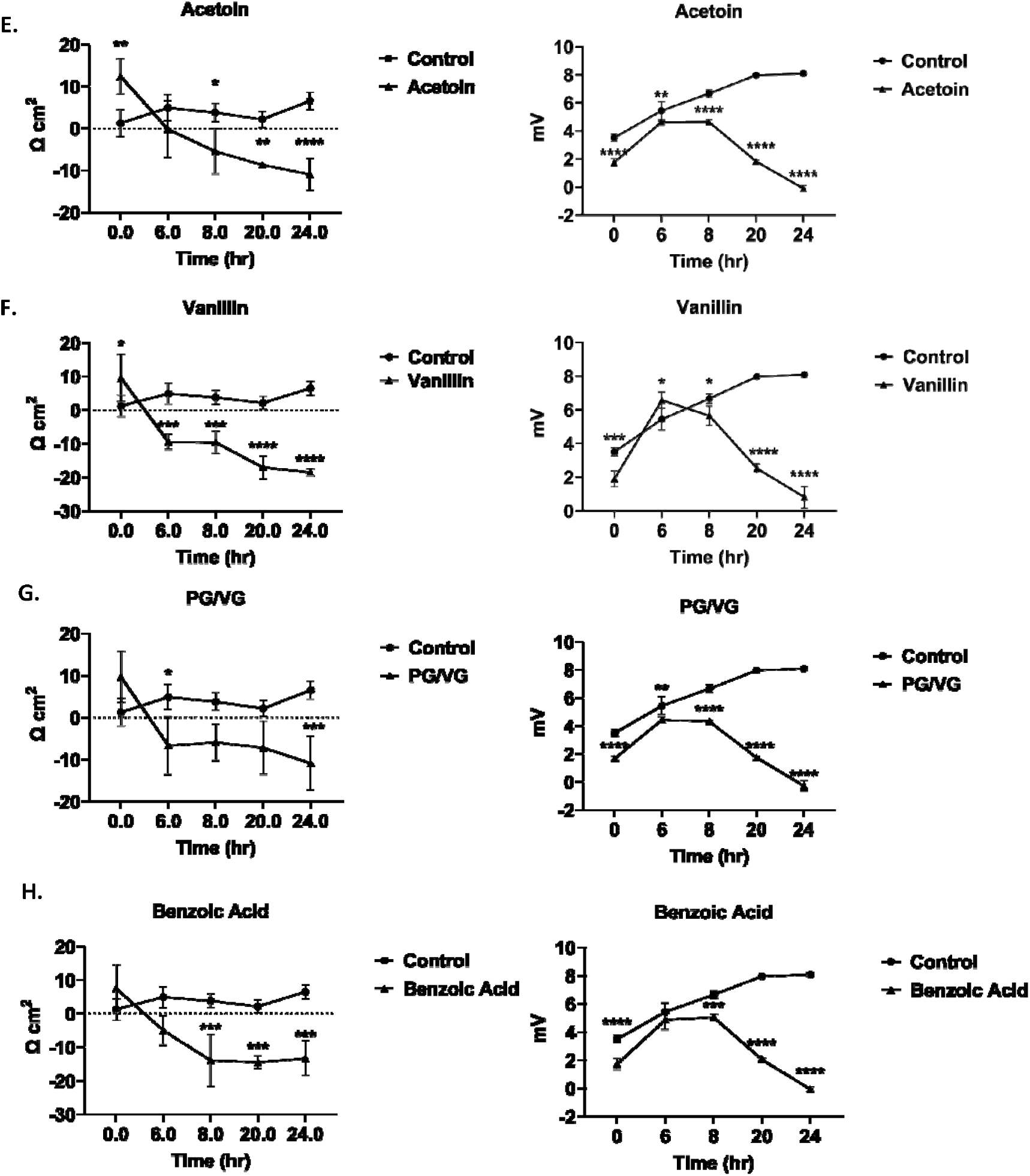
Menthol and tobacco flavoring chemicals caused BEAS-2B epithelial cell barrier dysfunction. BEAS2B cells were grown in transwell inserts in complete medium. Once reached a monolayer and full confluency, cells were serum deprived overnight. Cells were treated with 100 uM (A) 98% Menthone. (B) L-Menthone, (C). Carvone (D) WS-23 (E) Acetoin, (F) Vanillin, (G) PG/VG, and (H) Benzoic Acid. Transepithelial electrical resistance (TEER) and voltage (mV) data were collected pretreatment (0 hr), 6, 8, 20, and 24 hrs. following the treatments and the linear regression of TEER and mV ± SEM are represented. *p<0.05, **p<0.01, ***p<0.001, ****p<0.0001 vs untreated control., one-way ANOVA. N=3 wells per chemical treatment.

### ENDS constituents elicited an inflammatory response in bronchial epithelial cells

Inflammatory mediators, IL6 and IL8, levels were measured in conditioned media to assess the elicited inflammatory response in lung epithelial cells by commonly found ENDS constituents in menthol and tobacco flavored products **(Fig. 2)**. PG/VG caused a significant IL6 increase **(Fig. 2A)**. IL6 levels were increased in L-menthone, 98% menthone, vanillin, and acetoin treated groups compared to the untreated controls **(Fig. 2B)**. Carvone and WS-23 elicited a significant response (*p<0.05) (**Fig. 2B)**.

**Figure 2:**
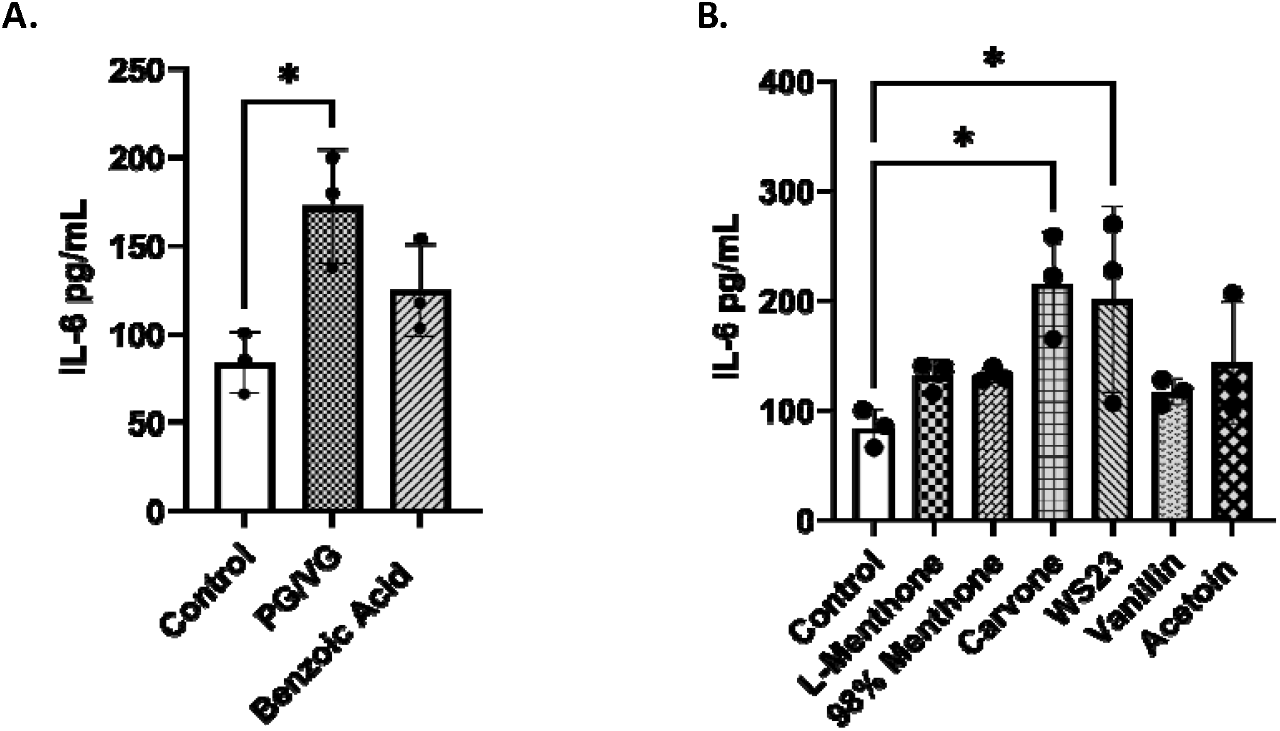
Menthol and tobacco flavoring constituents elicited an interleukin 6 cytokine response in lung epithelial cells. BEAS-2B cells cultured in transwells in complete media, at full confluency serum deprived overnight. Cells were treated with 100uM L-menthone, 98% menthone, carvone, WS-23, vanillin, acetoin, benzoic acid, and PG/VG. Apical conditioned media was collected after the 24-hour time point and IL6 was quantified. (A) control, PG/VG, and benzoic acid-induced response, and (B) L-menthone, 98% menthone, carvone, WS-23, vanillin, and acetoin response compared to untreated control. IL6 concentration in pg/mL ± SEM is represented, *p<0.05. vs. control, one-way ANOVA. N=3 wells per treatment

Pro-inflammatory cytokine IL-8 was significantly increased by 98% menthone (*p<0.05) and carvone (**p<0.05) compared to the untreated control cells **(Fig. 3)**

**Figure 3:**
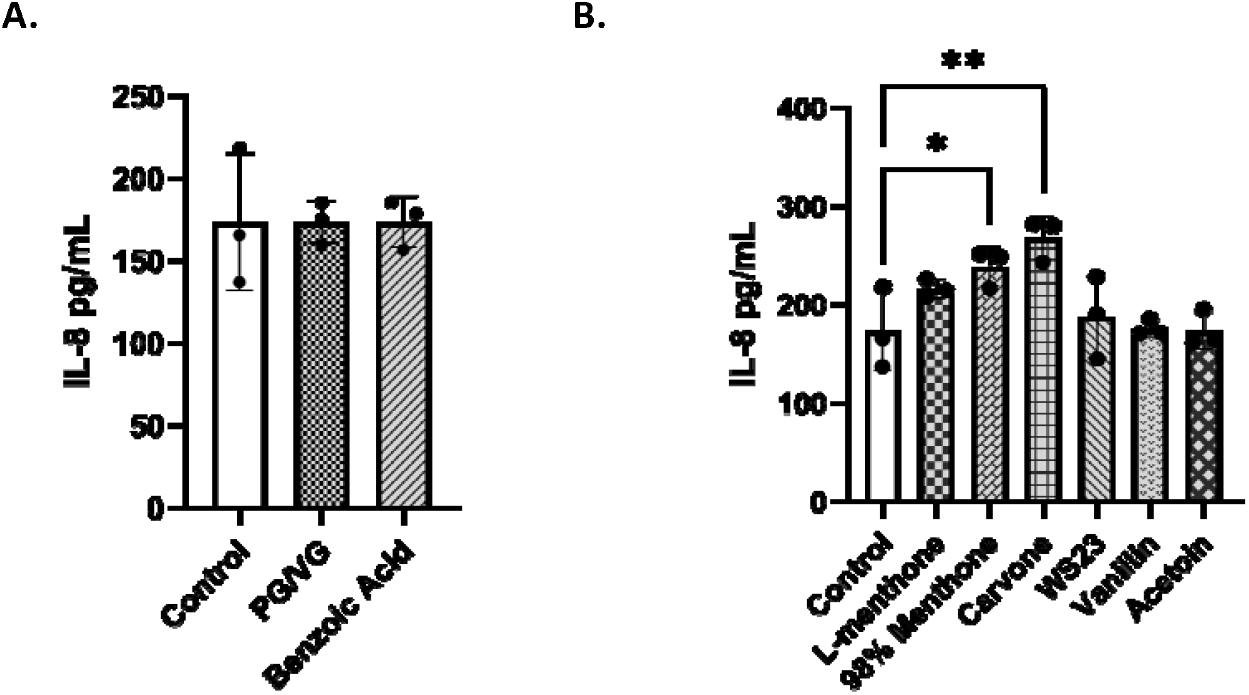
Menthol and tobacco flavoring constituents elicited an interleukin-8 cytokine response in lung epithelial cells. BEAS-2B cells cultured in transwells in complete media, at full confluency serum deprived overnight. Cells were treated with 100 uM L-menthone, 98% menthone, carvone, WS-23, vanillin, acetoin, benzoic acid, and PG/VG. Apical conditioned media was collected after the 24-hour time point and IL6 was quantified. (A) control, PG/VG, and benzoic acid-induced response, and (B) L-menthone, 98% menthone, carvone, WS-23, vanillin, and acetoin response compared to untreated control. IL8 concentration in pg/mL ± SEM is represented, *p<0.05, and **p<0.01 vs. untreated control. one-way ANOVA. N=3 wells per treatment

### Cytotoxicity Assessment

BEAS-2B cells were collected after the 24-hour point following the treatments and the cytotoxicity was determined by AO/PI staining. Compared to the untreated control, the cells treated with most constituents were showed negligible cytotoxicity (<10% cell death) **(Fig. 4)**. The highest toxicity was observed in cells treated with 98% Menthone (∼19%) (*p<0.05). This suggests that for the tested chemicals, PG/VG, benzoic acid, carvone, WS23, vanillin, and acetoin, the tested concentration (100 uM) was sufficient to induce the observed responses without causing cell death.

**Figure 4:**
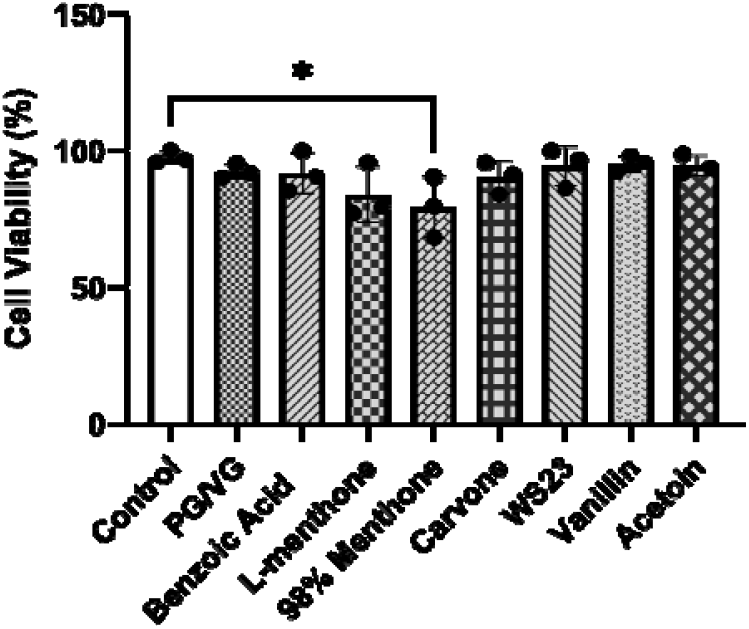
Menthol and tobacco flavoring constituents caused minimum cytotoxicity BEAS-2B cells. BEAS-2B cells cultured in transwells in complete media, at full confluency serum deprived overnight. Cells were treated with 100 uM L-menthone, 98% menthone, carvone, WS-23, vanillin, acetoin, benzoic acid, and PG/VG. At the 24 hr time point, cells were collected and stained with acridine orange and propidium iodide and the live, cell, and total cells were counted using CellDrop automatic cell counter. Cytotoxicity±SEM is represented. *p<0.05 vs. control, one-way ANOVA, N=3 wells per treatment.

### Non-nicotine Flavoring chemicals increased nicotinic acetylcholine receptors in bronchial epithelial cells

Prepared lysates of menthol and tobacco constituent treated cells were used to perform immunoblotting to analyze protein abundance of nicotinic acetylcholine receptors 1-5(CHRNAs) compared to untreated controls. The data was normalized via housekeeping protein beta-actin **(Fig. 5)**. CHRNA1 was increased by acetoin and PG/VG treatments (not significant) **(Fig 5A)**. CHRNA 4 was increased by carvone and acetoin (not significant) **(Fig 5B)**. A significant increase in the CHRNA5 for cells treated with acetoin and PG/VG compared to the control group (p****<0.0001) while menthone and 98% menthol showed no change **(Fig. 5C, D)**. Additionally, a significant increase in protein abundance for CHRNA7 was observed in cells treated with WS-23 (*p<0.05) but carvone did not cause any changes. These data suggest that CHRNA 5 and 7 were more affected compared to other CHRNAs by the tested chemicals.

**Figure 5:**
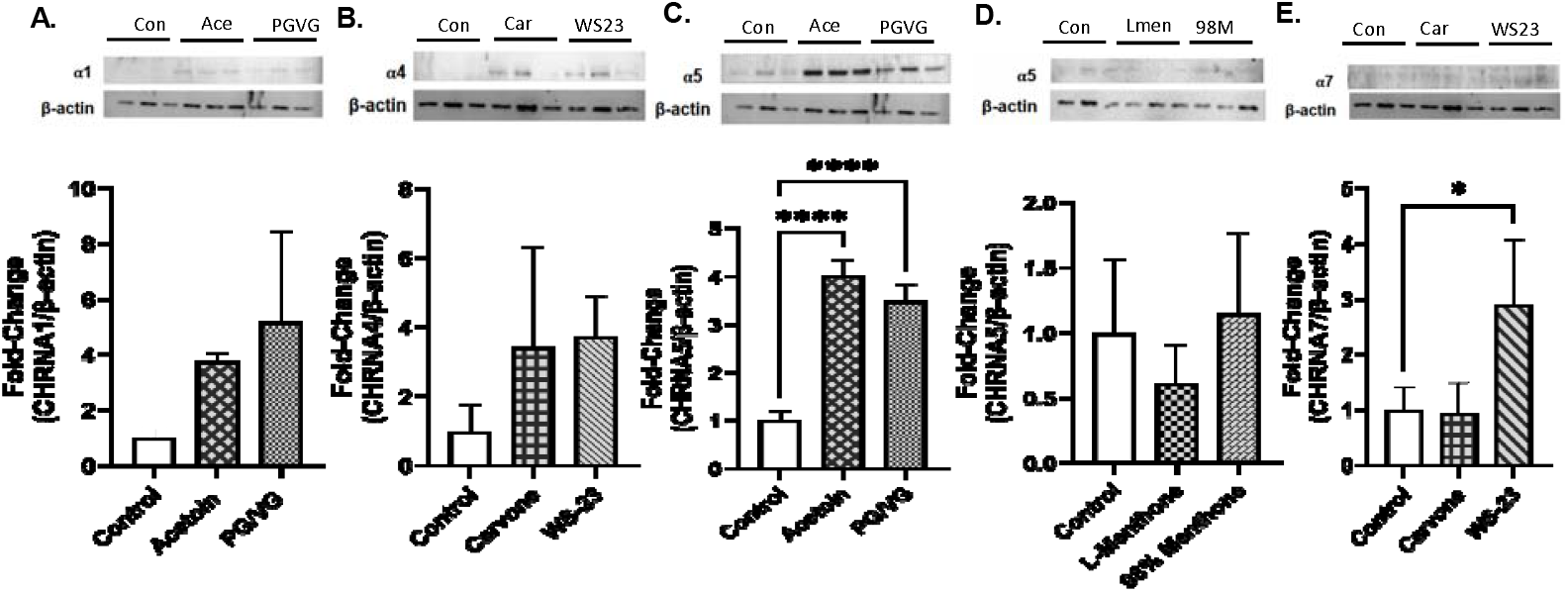
Menthol and tobacco flavoring constituents caused nicotinic acetylcholine receptor (CHRNA) modulation in BEAS2B lung epithelial cells. BEAS-2B cells cultured in transwells in complete media, at full confluency serum deprived overnight. Cells were treated with 100 uM L-menthone, 98% menthone, carvone, WS-23, vanillin, acetoin, benzoic acid, and PG/VG. At the 24 hr time point, cells were collected, lysed, and after BCA protein estimation, 5-10 ug of protein were loaded to 10-well gel for SDS-gel electrophoresis. After cellulose membrane transfer and blocking, the membranes were probed with primary antibodies for CHRNA 1,4,5, and 7, with *β*-actin loading control for normalization. Same membrane was sometimes re-probed up to 3 times with a different CHRNA. The blots with (A) Nicotinic Acetylcholine Receptors Alpha 1 expression with Acetoin and PG/VG. (B) Nicotinic Acetylcholine Receptors Alpha 4 expression with Carvone and WS-23. (C) Nicotinic Acetylcholine Receptors Alpha 5 expression with Acetoin and PG/VG. (D) Nicotinic Acetylcholine Receptors Alpha 5 expression with L-Menthone and 98% Menthone. (E) Nicotinic Acetylcholine Receptors Alpha 7 expression with Carvone and WS-23. All respective CHRNA bands *β* actin are shown with their densitometry fold-change ± SEM. *p<0.05 and ****p<0.0001 vs. control, one-way ANOVA. N=3 wells per chemical. Full blots are shown in the supplementary section.

## Discussion

Electronic cigarettes are becoming widely popular and menthol and tobacco flavors are used as a safer alternative for adult users and a cessation product to combustible cigarettes(5, 26). While these flavors constitute numerous chemicals, key chemicals that are commonly used in menthol- and tobacco-flavored ENDS products such as acetoin, vanillin, 98% menthone, WS23, carvone, and base liquid constituent PG/VG and benzoic acid were selected to test the hypothesis that low concentrations in liquids can cause lung epithelial barrier function, elicit an inflammatory response, and modulate nicotinic acetylcholine receptors independent of nicotine. Similar to our study, these chemicals have been used due to their prevalence in several studies involving the characterization of E-liquids, specifically, the ones that were menthol and tobacco flavored **(Table 1)**.

Our data demonstrated acute adverse effects on epithelial barrier function by decreasing TEER and mV values. While, acetoin, vanillin, WS23, and PG/VG caused a significant decrease in both TEER and mV across the membrane, suggesting the disruption of the tight junctions by the chemicals, L-menthone, 98% menthone, and carvone caused a severe decline in voltage across the membrane suggesting, even though the tight junctions were not impaired or leaking, epithelial polarity, metabolism, and ion transport have been adversely affected as before structural breakdown(27, 28). These findings corroborate with studies where ENDS aerosols have been shown to cause barrier dysfunction (29, 30).

Our cytotoxicity data demonstrated the concentration we tested in the study at 100 uM was sufficient to induce a response but did not cause significant cell death, except for 98% menthone. Consistent with these results, other studies have shown that menthol flavors have greater toxicity in their BEAS2B 16HBE and CALU3 upon ENDS chemical and aerosol exposures (29-32).

Our data showed increased IL6 and IL8 pro-inflammatory cytokine response by PG/VG, carvone, WS-23, L-menthone, and 98% menthone. These cytokines initiate and amplify the inflammatory response by recruiting other immune cells, such as neutrophils, in the acute phase. These cytokines are often increased in acute lung injury, asthma, and infections(14, 33). Even though we saw a significant cell death by 98%, we observed significantly elevated cytokine levels, suggesting the cell death occurred around the 20 hr time point based on declined TEER and mV drop around the same time.

Acetoin has also been shown to be a precursor for diacetyl (22) inhalation of it has been linked with bronchiolitis obliterans in many flavoring manufacturing workers (34). Benzoic acid, often used for nicotine salts chemical exposers from previous literature (35) showed a decrease in the overall TEER results from a 24-hour exposure of 3% PG/VG in Human bronchiole epithelial cells. Consistent with our cytokine results, exposure to aerosolized PG/VG in normal human bronchial epithelial cells (HBEC) cells showed increased mRNA markers for both IL-6 and IL-8, further supporting our findings (36). These studies corroborate the adverse cellular effects we observed by benzoic acid and PG/VG, such as barrier dysfunction and inflammation.

In our study, WS23, a synthetic cooling agent added to vaping products, showed no less comparative toxicity to other menthol analogs. We observed barrier dysfunction, IL6 elevation, and CHRNA7 increase by WS23. Our data and similar studies on airway epithelial cells with aerosolized WS-23 and nicotine showed an increased IL-6 following a 72-hour exposure, suggesting that synthetic cooling agents induce lung inflammation (37). Our study demonstrated that WS23 containing “ice” vaping products may pose similar pulmonary toxicity in comparison to other menthol additives (38).

Nicotinic acetylcholine receptors (CHRNAs) are cholinergic and are typically activated by binding to their ligands, nicotine and acetylcholine, and are known to be involved in lung disease pathogenesis(6, 9, 39, 40). In BEAS2B lysates, we determined the CHRNA subunit dysregulated expression after treatment with the chemicals. Our data showed that PG/VG, acetoin, and WS23 significantly augmented the a5 and a7 subunits compared to untreated counterparts, suggesting receptor desensitization, immunomodulation, and increased risk for inflammatory diseases like COPD. Supporting our data, a7 receptor-mediated pathways have been shown to play a role in airway remodeling and obstruction, contributing to the progression of COPD (6). Additionally, it also mediated pathways responsible for lung cancer in the presence of nicotine (41). These subunits are involved in JAK/STAT, ERK1/2, and PI3K/Akt mechanisms of metabolism, proliferation, and inflammation(40). We also demonstrated increased a1 and a4 involved by PG/VG, acetoin, carvone, and WS23, which may suggest stress response and cytokine modulation. CHRNA1(a1), neuromuscular receptor increases may also be due to abnormal receptor remodeling. Overall, increased CHRNA expression demonstrated the increased sensitivity to these non-nicotinic chemicals, potentially mediating lung diseases similar to nicotine receptor signaling. This indicates the possible pathogenesis of pulmonary diseases such as emphysema/COPD with e-cigarette use, even in the absence of nicotine. Further investigation is required to learn more about the mechanism of CHRNA receptor signaling in lung disease pathogenesis by non-nicotinic receptors.

In summary, our data showed that commonly used non-nicotine-constituents of menthol and tobacco-flavored ENDS modulate nicotinic receptor expression on lung epithelial cells, cause lung epithelial barrier dysfunction–by altering polarization, hindering ion transport, and by disrupting the tight-junctions–and elicit cytokine responses such as IL6 and IL8 which are involved in innate immunity and acute lung injury responses.

## Conclusion

Our data support the hypothesis that chemicals in e-liquids cause epithelial barrier dysfunction, induce lung inflammation, and alter nicotinic receptor signaling pathways, leading to lung diseases. The findings provide insights into regulating constituents and their concentrations to achieve potentially safer levels.

## Supporting information

Supplementary Figures- Western Full Blots

## List of Abbreviations

COPD: Chronic obstructive pulmonary disease
ELISA: Enzyme-linked immunosorbent assay
PG/VG: Propylene glycol/Vegetable glycerol
AO/PI: Acridine orange/Propidium iodide
IL-6: Interleukin-6
IL-8: Interleukin-8
ENDS: Electronic nicotine delivery systems
PATH: Population Assessment of Tobacco and Health
ARDS: Acute respiratory distress syndrome
TNF: Tumor necrosis factor
BALF: Bronchoalveolar lavage fluid
ALI: Acute lung injury
ALI: Air-liquid interphase
NF-κB: Nuclear factor kappa B
FBS: Fetal bovine serum
HEPES: 4-(2-hydroxyethyl)piperazine-1-ethanesulfonic acid
DMEM: Dulbecco’s Modified Eagle Medium
TEER: Transepithelial electrical resistance
EDTA: Ethylenediaminetetraacetic acid
HBEC: Primary human bronchial epithelial cells
CHRNA: Nicotinic Acetylcholine Receptors

## Availability of data and materials

All data generated or analyzed during this study are included in this published article and raw data available upon request.

## Authors’ contributions

TM conceptualized and designed the experiments. VSP. KJB, and AB performed the experiments. VSP and AB wrote the manuscript. TM and KJB edited the manuscript.

## Acknowledgements

Purdue PURE Tox summer program for supporting Vidhi Pandya

## References

1. Kaplan B, Hardesty JJ, Welding K, Breland AB, Eissenberg T, Cohen JE. Electronic Nicotine Delivery System flavor use over time by age group in the US: A longitudinal analysis. Tob Induc Dis. 2023;21:67.

2. Sharma E, Zebrak K, Lauten K, Gravely S, Cooper M, Gardner LD, et al. Cigarette and ENDS dual use longitudinal transitions among adults in the Population Assessment of Tobacco and Health (PATH) Study, Waves 4-5 (2016-2019). Addict Behav Rep. 2024;19:100528.

3. Romm KF, Henriksen L, Huang J, Le D, Clausen M, Duan Z, et al. Impact of existing and potential e-cigarette flavor restrictions on e-cigarette use among young adult e-cigarette users in 6 US metropolitan areas. Prev Med Rep. 2022;28:101901.

4. Wise PM, Breslin PA, Dalton P. Sweet taste and menthol increase cough reflex thresholds. Pulm Pharmacol Ther. 2012;25(3):236–41.

5. Rest EC, Brikmanis KN, Mermelstein RJ. Preferred flavors and tobacco use patterns in adult dual users of cigarettes and ENDS. Addict Behav. 2022;125:107168.

6. Hollenhorst MI, Krasteva-Christ G. Nicotinic Acetylcholine Receptors in the Respiratory Tract. Molecules. 2021;26(20).

7. Roman J, Koval M. Control of lung epithelial growth by a nicotinic acetylcholine receptor: the other side of the coin. Am J Pathol. 2009;175(5):1799–801.

8. Herman M, Tarran R. E-cigarettes, nicotine, the lung and the brain: multi-level cascading pathophysiology. J Physiol. 2020;598(22):5063–71.

9. Diabasana Z, Perotin JM, Belgacemi R, Ancel J, Mulette P, Delepine G, et al. Nicotinic Receptor Subunits Atlas in the Adult Human Lung. Int J Mol Sci. 2020;21(20).

10. King JR, Ullah A, Bak E, Jafri MS, Kabbani N. Ionotropic and Metabotropic Mechanisms of Allosteric Modulation of α7 Nicotinic Receptor Intracellular Calcium. Mol Pharmacol. 2018;93(6):601–11.

11. McElvaney OJ, Curley GF, Rose-John S, McElvaney NG. Interleukin-6: obstacles to targeting a complex cytokine in critical illness. Lancet Respir Med. 2021;9(6):643–54.

12. Barnes PJ. The cytokine network in asthma and chronic obstructive pulmonary disease. J Clin Invest. 2008;118(11):3546–56.

13. Qazi BS, Tang K, Qazi A. Recent advances in underlying pathologies provide insight into interleukin-8 expression-mediated inflammation and angiogenesis. Int J Inflam. 2011;2011:908468.

14. Rincon M, Irvin CG. Role of IL-6 in asthma and other inflammatory pulmonary diseases. Int J Biol Sci. 2012;8(9):1281–90.

15. Donnelly SC, Strieter RM, Kunkel SL, Walz A, Robertson CR, Carter DC, et al. Interleukin-8 and development of adult respiratory distress syndrome in at-risk patient groups. Lancet. 1993;341(8846):643–7.

16. Marini M, Vittori E, Hollemborg J, Mattoli S. Expression of the potent inflammatory cytokines, granulocyte-macrophage-colony-stimulating factor and interleukin-6 and interleukin-8, in bronchial epithelial cells of patients with asthma. J Allergy Clin Immunol. 1992;89(5):1001–9.

17. Tillie-Leblond I, Pugin J, Marquette CH, Lamblin C, Saulnier F, Brichet A, et al. Balance between pro-inflammatory cytokines and their inhibitors in bronchial lavage from patients with status asthmaticus. Am J Respir Crit Care Med. 1999;159(2):487–94.

18. Shute JK, Vrugt B, Lindley IJ, Holgate ST, Bron A, Aalbers R, et al. Free and complexed interleukin-8 in blood and bronchial mucosa in asthma. Am J Respir Crit Care Med. 1997;155(6):1877–83.

19. Hoffmann E, Dittrich-Breiholz O, Holtmann H, Kracht M. Multiple control of interleukin-8 gene expression. J Leukoc Biol. 2002;72(5):847–55.

20. Baughman RP, Gunther KL, Rashkin MC, Keeton DA, Pattishall EN. Changes in the inflammatory response of the lung during acute respiratory distress syndrome: prognostic indicators. Am J Respir Crit Care Med. 1996;154(1):76–81.

21. Norzila MZ, Fakes K, Henry RL, Simpson J, Gibson PG. Interleukin-8 secretion and neutrophil recruitment accompanies induced sputum eosinophil activation in children with acute asthma. Am J Respir Crit Care Med. 2000;161(3 Pt 1):769–74.

22. Vas CA, Porter A, McAdam K. Acetoin is a precursor to diacetyl in e-cigarette liquids. Food Chem Toxicol. 2019;133:110727.

23. Hsiao YC, Matulewicz RS, Sherman SE, Jaspers I, Weitzman ML, Gordon T, et al. Untargeted Metabolomics to Characterize the Urinary Chemical Landscape of E-Cigarette Users. Chem Res Toxicol. 2023;36(4):630–42.

24. Durham AL, Adcock IM. The relationship between COPD and lung cancer. Lung Cancer. 2015;90(2):121–7.

25. Pell TJ, Gray MB, Hopkins SJ, Kasprowicz R, Porter JD, Reeves T, et al. Epithelial Barrier Integrity Profiling: Combined Approach Using Cellular Junctional Complex Imaging and Transepithelial Electrical Resistance. SLAS Discov. 2021;26(7):909–21.

26. Joseph M, Morean ME, Wu R, Krishnan-Sarin S, O’Malley SS, Bold KW. Examining E-Cigarette Flavor Use and Preference by Menthol Cigarette Status and Quit Duration Among US Adults Using E-Cigarettes in a Smoking Cessation Attempt. Nicotine Tob Res. 2025.

27. Tilston-Lunel AM, Varelas X. Polarity in respiratory development, homeostasis and disease. Curr Top Dev Biol. 2023;154:285–315.

28. Vladar EK, Nayak JV, Milla CE, Axelrod JD. Airway epithelial homeostasis and planar cell polarity signaling depend on multiciliated cell differentiation. JCI Insight. 2016;1(13).

29. Muthumalage T, Lamb T, Friedman MR, Rahman I. E-cigarette flavored pods induce inflammation, epithelial barrier dysfunction, and DNA damage in lung epithelial cells and monocytes. Sci Rep. 2019;9(1):19035.

30. Lamb T, Muthumalage T, Rahman I. Pod-based menthol and tobacco flavored e-cigarettes cause mitochondrial dysfunction in lung epithelial cells. Toxicol Lett. 2020;333:303–11.

31. Rowell TR, Reeber SL, Lee SL, Harris RA, Nethery RC, Herring AH, et al. Flavored e-cigarette liquids reduce proliferation and viability in the CALU3 airway epithelial cell line. Am J Physiol Lung Cell Mol Physiol. 2017;313(1): L52–l66.

32. Muthumalage T, Prinz M, Ansah KO, Gerloff J, Sundar IK, Rahman I. Inflammatory and Oxidative Responses Induced by Exposure to Commonly Used e-Cigarette Flavoring Chemicals and Flavored e-Liquids without nicotine. Front Physiol. 2017;8:1130.

33. Cesta MC, Zippoli M, Marsiglia C, Gavioli EM, Mantelli F, Allegretti M, et al. The Role of Interleukin-8 in Lung Inflammation and Injury: Implications for the Management of COVID-19 and Hyperinflammatory Acute Respiratory Distress Syndrome. Front Pharmacol. 2021;12:808797.

34. Kreiss K, Fedan KB, Nasrullah M, Kim TJ, Materna BL, Prudhomme JC, et al. Longitudinal lung function declines among California flavoring manufacturing workers. Am J Ind Med. 2012;55(8):657–68.

35. Woodall M, Jacob J, Kalsi KK, Schroeder V, Davis E, Kenyon B, et al. E-cigarette constituents propylene glycol and vegetable glycerin decrease glucose uptake and its metabolism in airway epithelial cells in vitro. Am J Physiol Lung Cell Mol Physiol. 2020;319(6): L957–l67.

36. Kim MD, Chung S, Baumlin N, Qian J, Montgomery RN, Sabater J, et al. The combination of propylene glycol and vegetable glycerin e-cigarette aerosols induces airway inflammation and mucus hyperconcentration. Sci Rep. 2024;14(1):1942.

37. Manevski M, Yogeswaran S, Rahman I, Devadoss D, Chand HS. Corrigendum to “E-cigarette synthetic cooling agent WS-23 and nicotine aerosols differentially modulate airway epithelial cell responses” [Toxicol. Rep. 9 (2022) 1823-1830]. Toxicol Rep. 2023;11:259–60.

38. Leventhal AM, Tackett AP, Whitted L, Jordt SE, Jabba SV. Ice flavours and non-menthol synthetic cooling agents in e-cigarette products: a review. Tob Control. 2023;32(6):769–77.

39. Lam DC, Luo SY, Fu KH, Lui MM, Chan KH, Wistuba, II, et al. Nicotinic acetylcholine receptor expression in human airway correlates with lung function. Am J Physiol Lung Cell Mol Physiol. 2016;310(3): L232–9.

40. Borkar NA, Thompson MA, Bartman CM, Khalfaoui L, Sine S, Sathish V, et al. Nicotinic receptors in airway disease. Am J Physiol Lung Cell Mol Physiol. 2024;326(2): L149–L63.

41. He Z, Xu Y, Rao Z, Zhang Z, Zhou J, Zhou T, et al. The role of α7-nAChR-mediated PI3K/AKT pathway in lung cancer induced by nicotine. Sci Total Environ. 2024;912:169604.

42. McAdam K, Waters G, Moldoveanu S, Margham J, Cunningham A, Vas C, et al. Diacetyl and Other Ketones in e-Cigarette Aerosols: Some Important Sources and Contributing Factors. Front Chem. 2021;9:742538.

43. Cardenas RB, Watson C, Valentin-Blasini L. Determination of Benzoic Acid and Benzyl Alcohol in E-Liquids (JUUL(™) Pods) by Isotopic Dilution High-Performance Liquid Chromatography and Tandem Mass Spectrometry. Contrib Tob Nicotine Res. 2021;30(4):212–20.

44. Tran LN, Rao G, Robertson NE, Hunsaker HC, Chiu EY, Poulin BA, et al. Quantification of Free Radicals from Vaping Electronic Cigarettes Containing Nicotine Salt Solutions with Different Organic Acid Types and Concentrations. Chem Res Toxicol. 2024;37(6):991–9.

45. Jabba SV, Erythropel HC, Torres DG, Delgado LA, Woodrow JG, Anastas PT, et al. Synthetic Cooling Agents in US-marketed E-cigarette Refill Liquids and Popular Disposable E-cigarettes: Chemical Analysis and Risk Assessment. Nicotine Tob Res. 2022;24(7):1037–46.

46. Tierney PA, Karpinski CD, Brown JE, Luo W, Pankow JF. Flavour chemicals in electronic cigarette fluids. Tob Control. 2016;25(e1):e10–5.

47. Effah F, Taiwo B, Baines D, Bailey A, Marczylo T. Pulmonary effects of e-liquid flavors: a systematic review. J Toxicol Environ Health B Crit Rev. 2022;25(7):343–71.

48. Kubica P. Determination of Glycerol, Propylene Glycol, and Nicotine as the Main Components in Refill Liquids for Electronic Cigarettes. Molecules. 2023;28(11).

49. Krüsemann EJZ, Pennings JLA, Cremers J, Bakker F, Boesveldt S, Talhout R. GC-MS analysis of e-cigarette refill solutions: A comparison of flavoring composition between flavor categories. J Pharm Biomed Anal. 2020;188:113364.

50. Omaiye EE, Luo W, McWhirter KJ, Pankow JF, Talbot P. Flavour chemicals, synthetic coolants and pulegone in popular mint-flavoured and menthol-flavoured e-cigarettes. Tob Control. 2022;31(e1):e3–e9.

51. Wang Q, Khan NA, Muthumalage T, Lawyer GR, McDonough SR, Chuang TD, et al. Dysregulated repair and inflammatory responses by e-cigarette-derived inhaled nicotine and humectant propylene glycol in a sex-dependent manner in mouse lung. FASEB Bioadv. 2019;1(10):609–23.

