## Supplementary Figures- Western Full Blots for "Comparative toxicity of menthol- and tobacco-flavored electronic cigarette constituents causing inflammation, epithelial barrier dysfunction, and nicotinic acetylcholine receptor modulation in the absence of nicotine"

### Slide 1
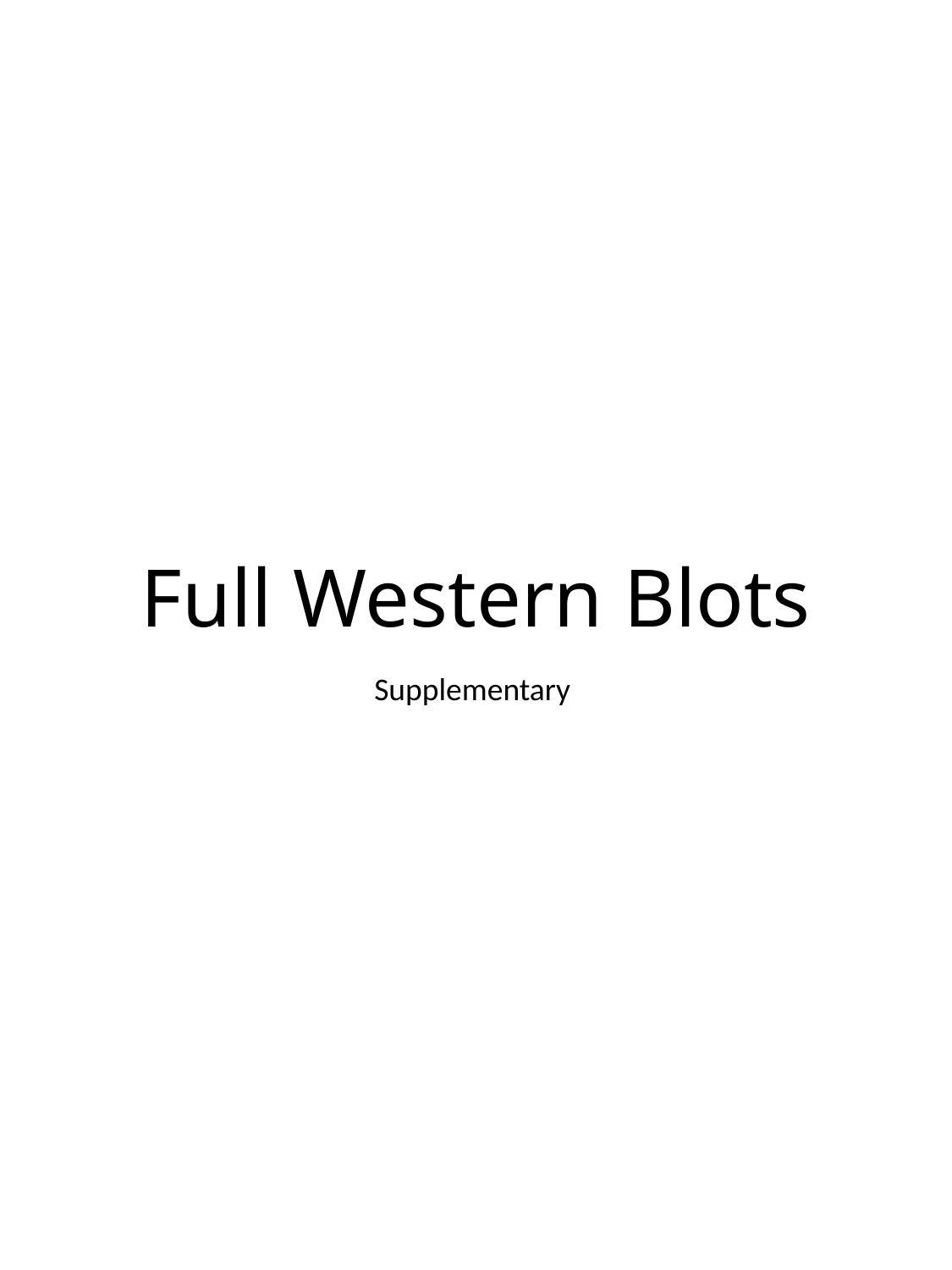

#### Full Western Blots
Supplementary

### Slide 2
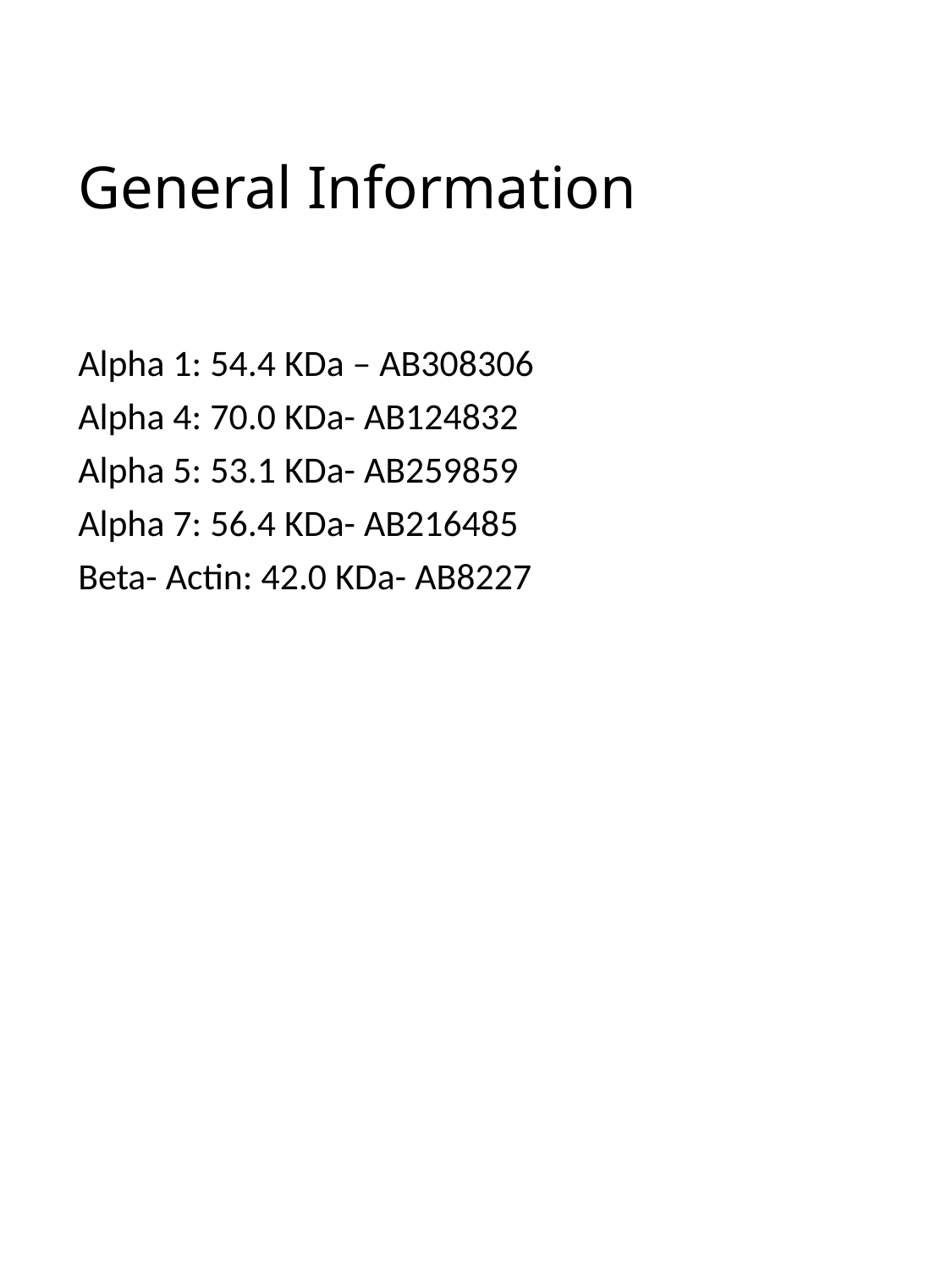

#### General Information
Alpha 1: 54.4 KDa – AB308306
Alpha 4: 70.0 KDa- AB124832
Alpha 5: 53.1 KDa- AB259859
Alpha 7: 56.4 KDa- AB216485
Beta- Actin: 42.0 KDa- AB8227

### Slide 3
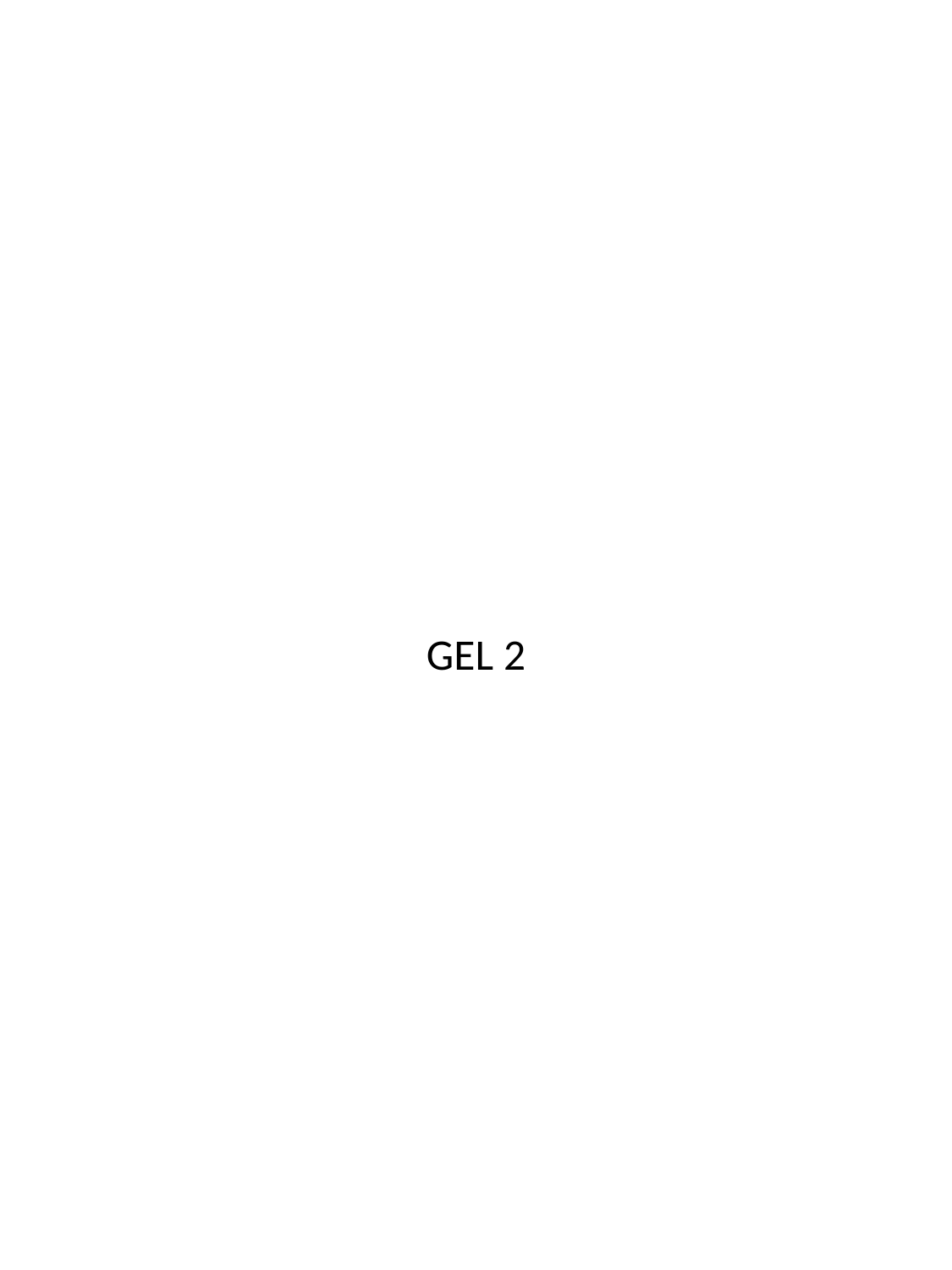

GEL 2

### Slide 4
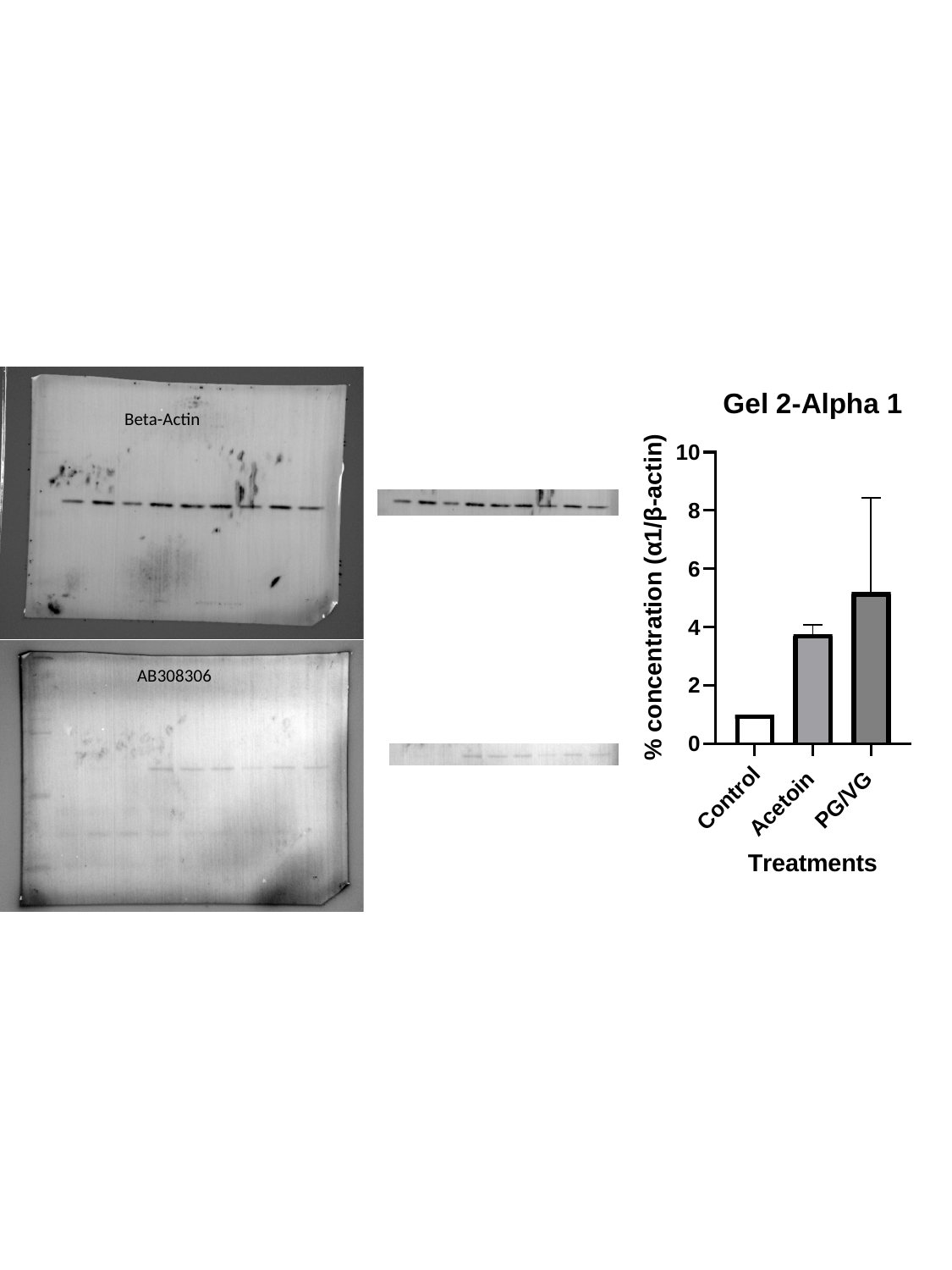

Beta-Actin
AB308306

### Slide 5
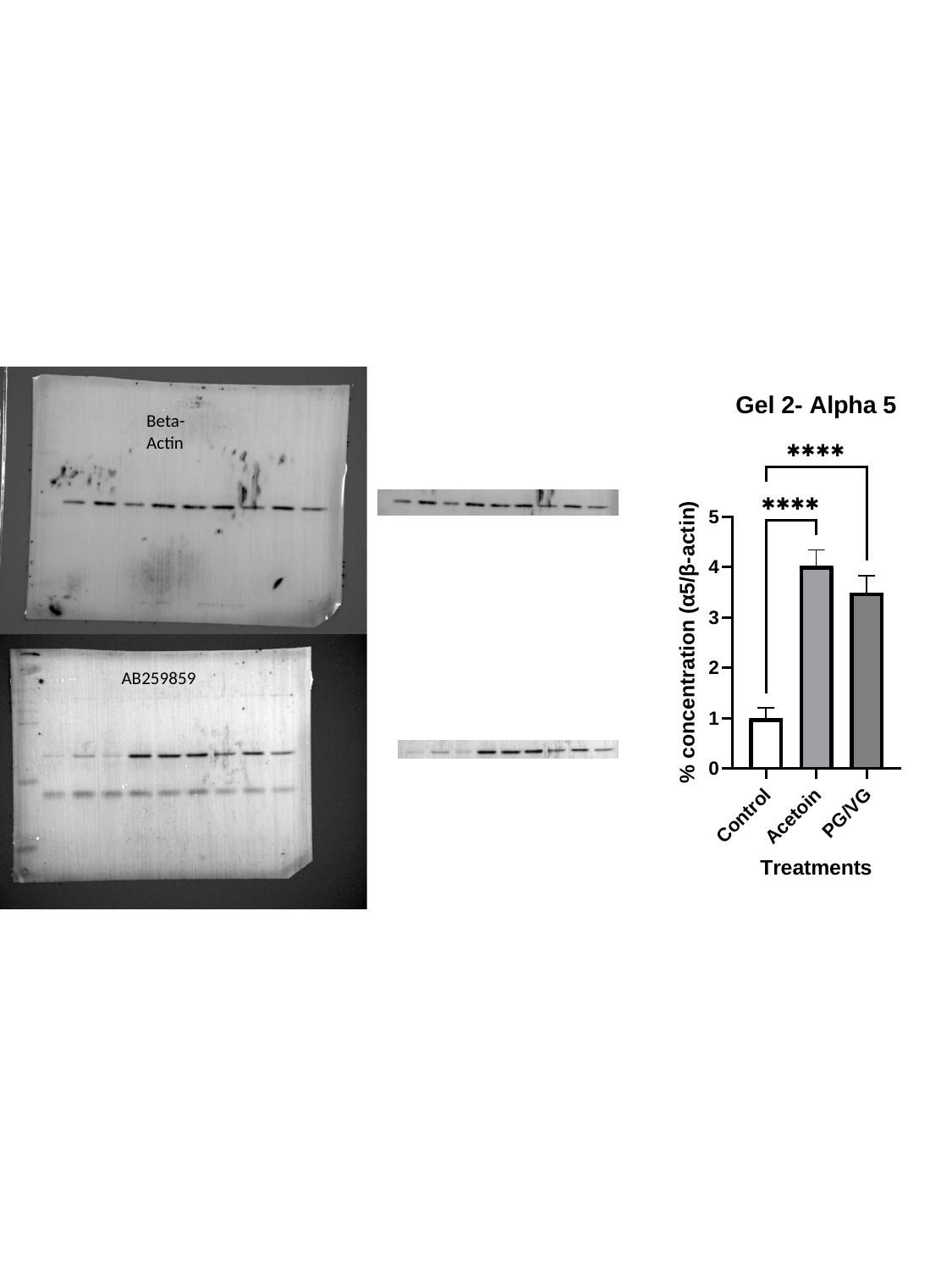

Beta-Actin
AB259859

### Slide 6
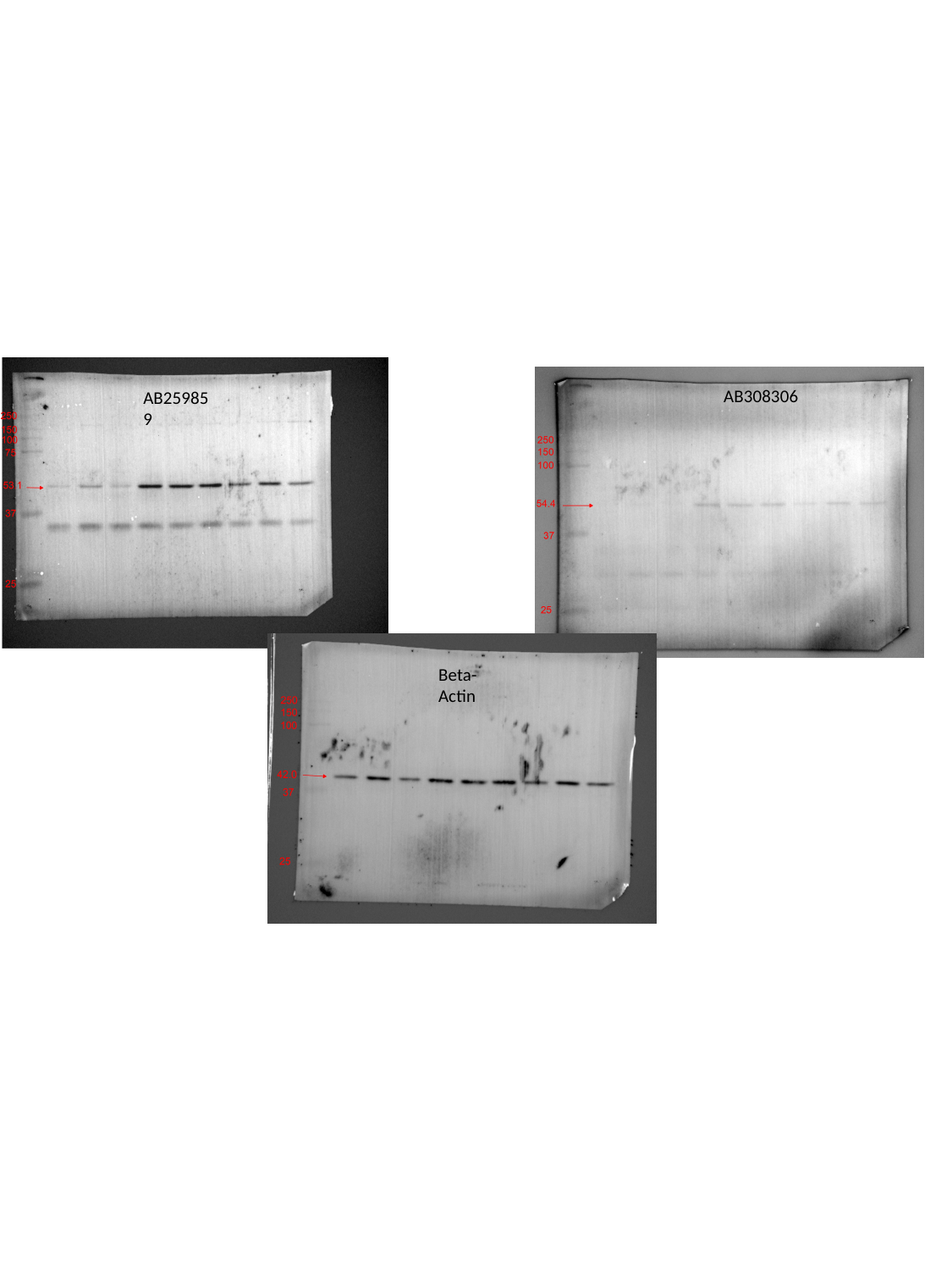

AB308306
AB259859
Beta-Actin

### Slide 7
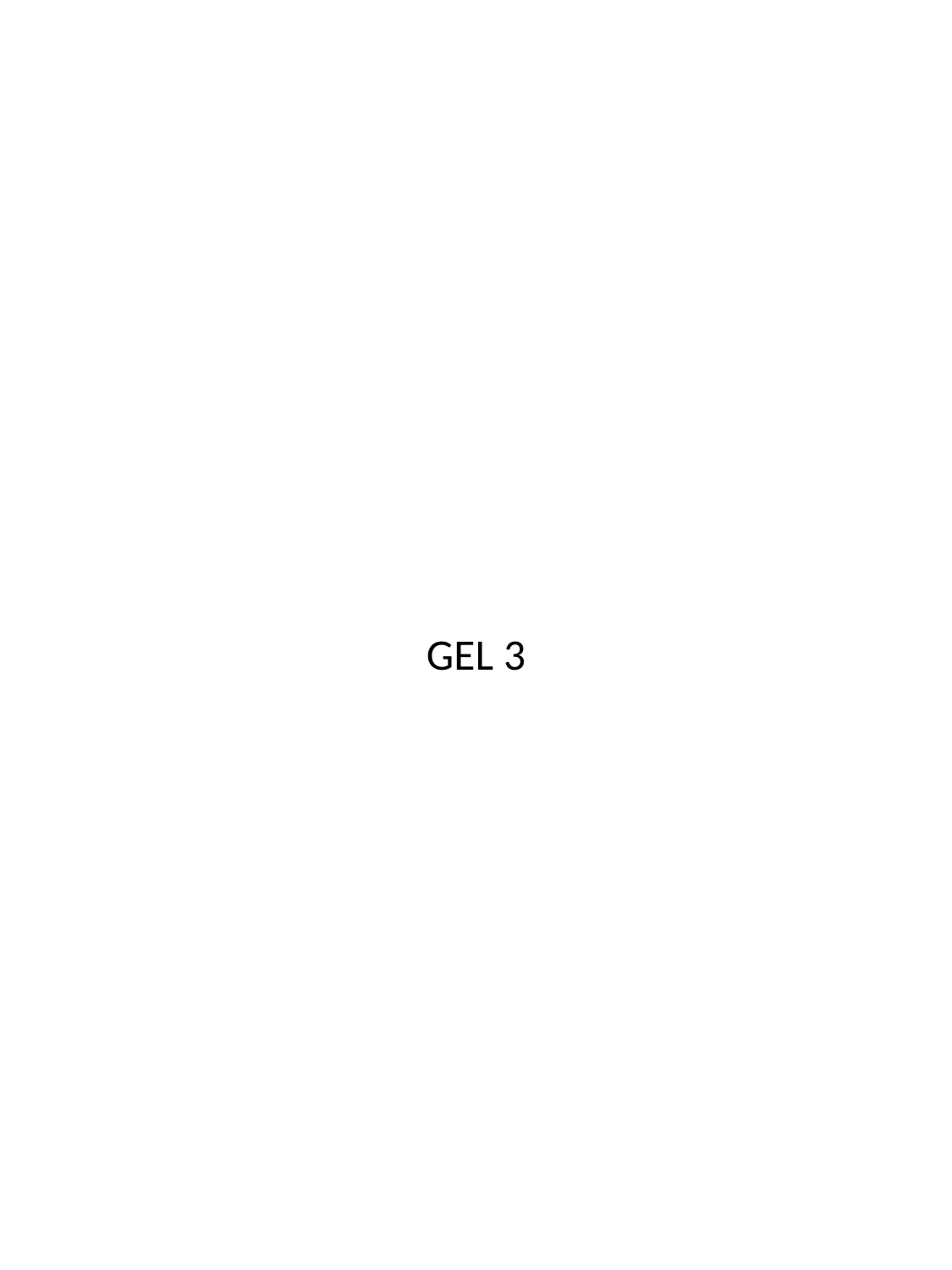

GEL 3

### Slide 8
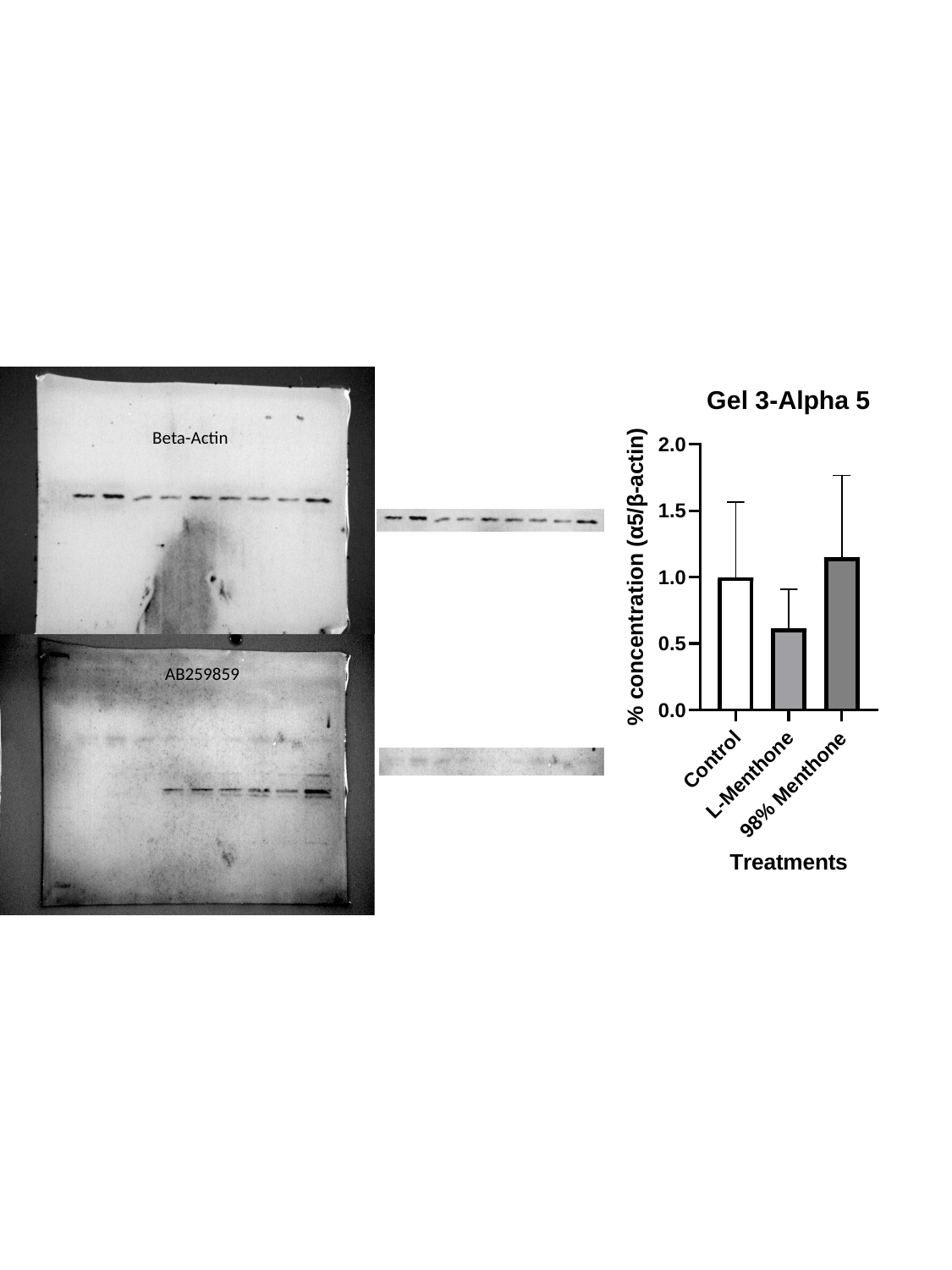

Beta-Actin
AB259859

### Slide 9
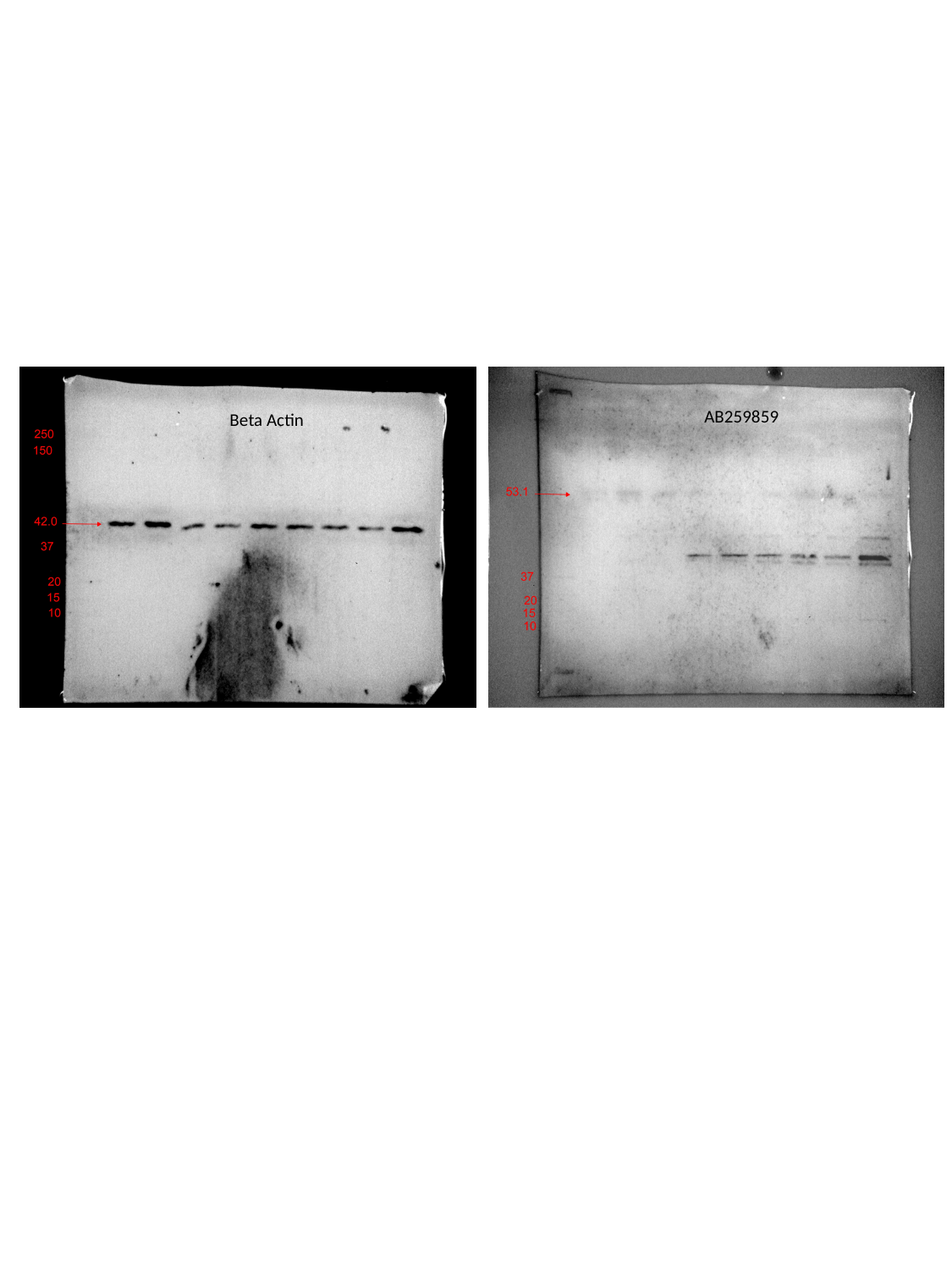

AB259859
Beta Actin

### Slide 10
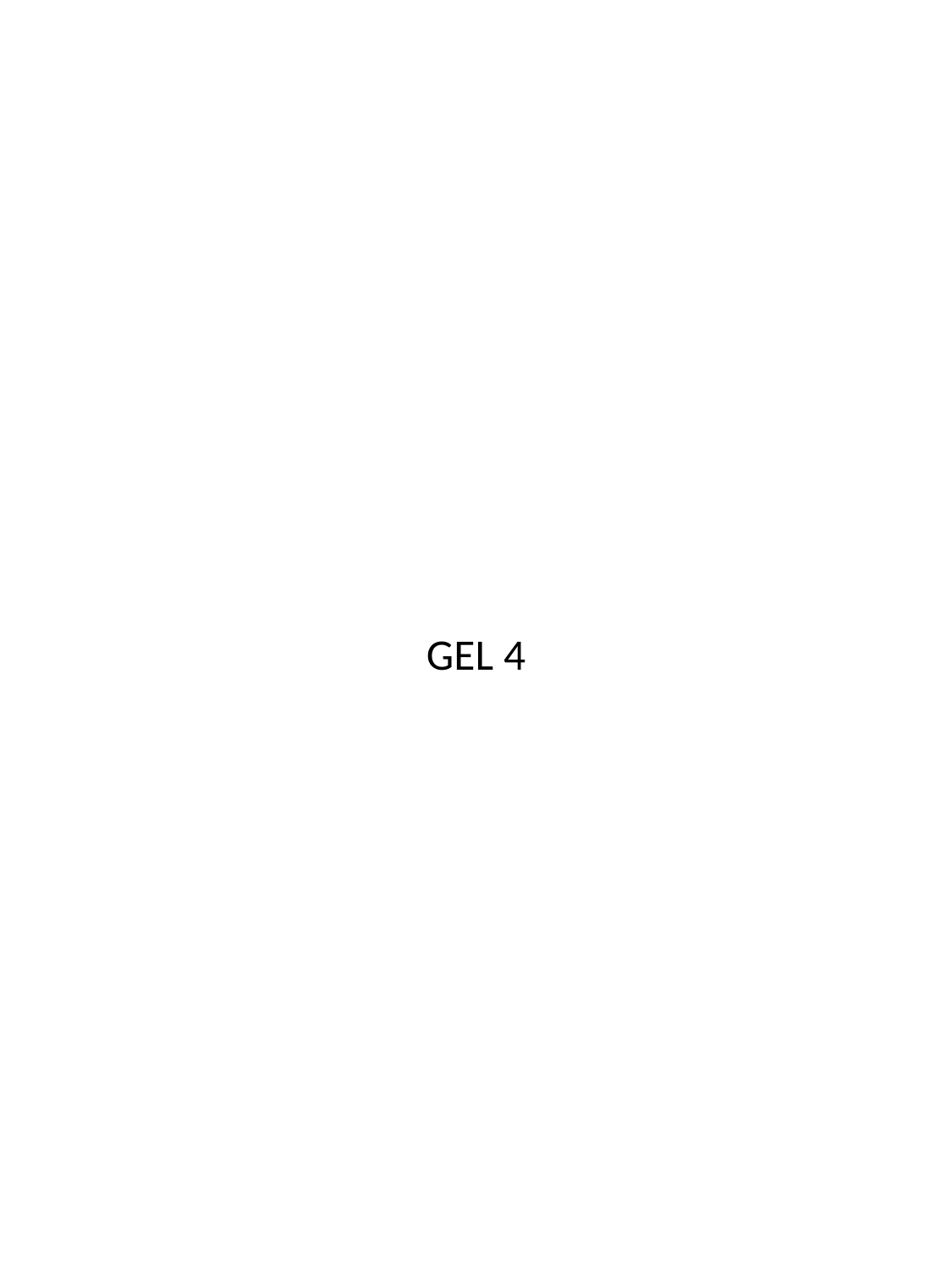

GEL 4

### Slide 11
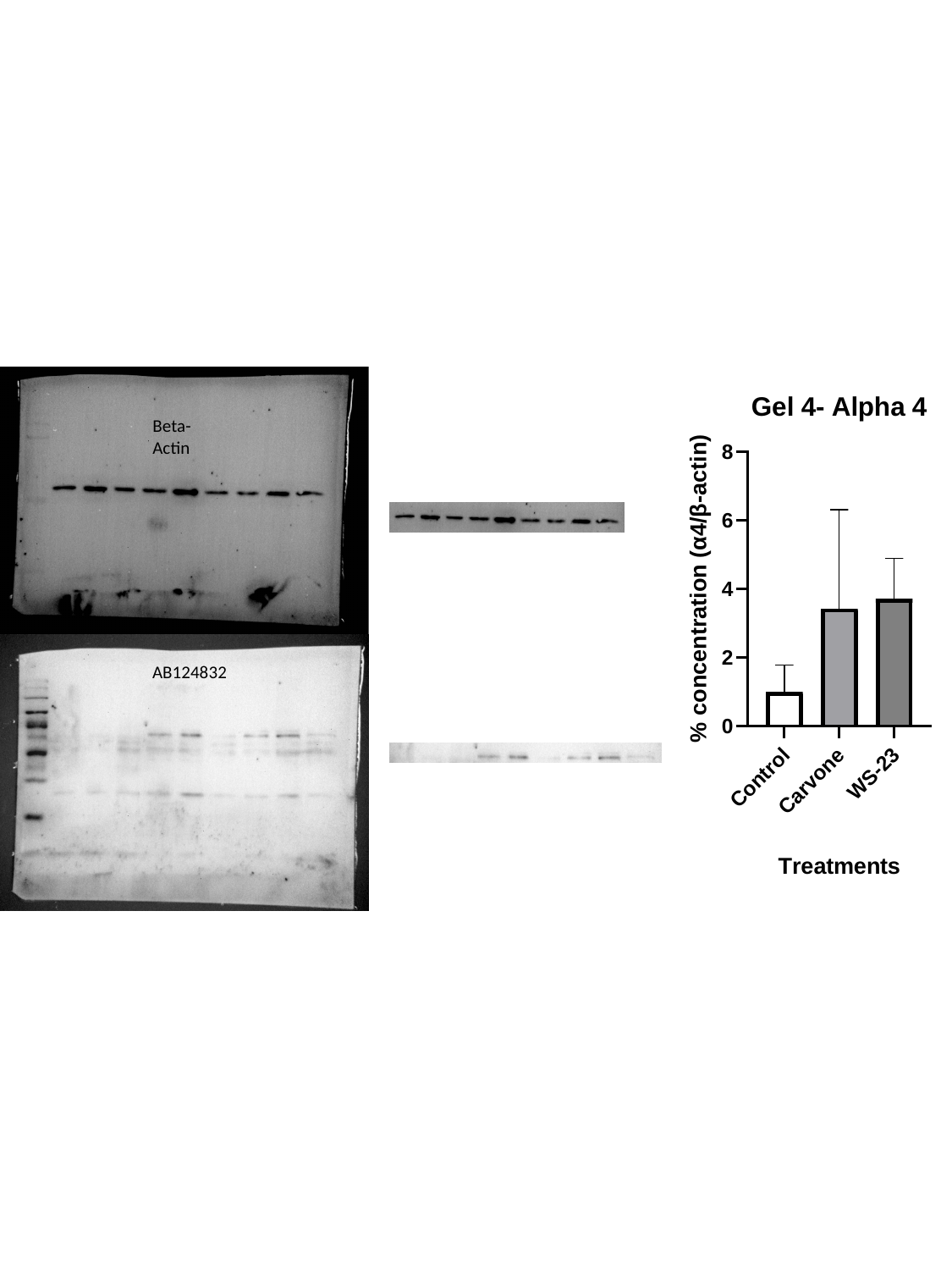

Beta-Actin
AB124832

### Slide 12
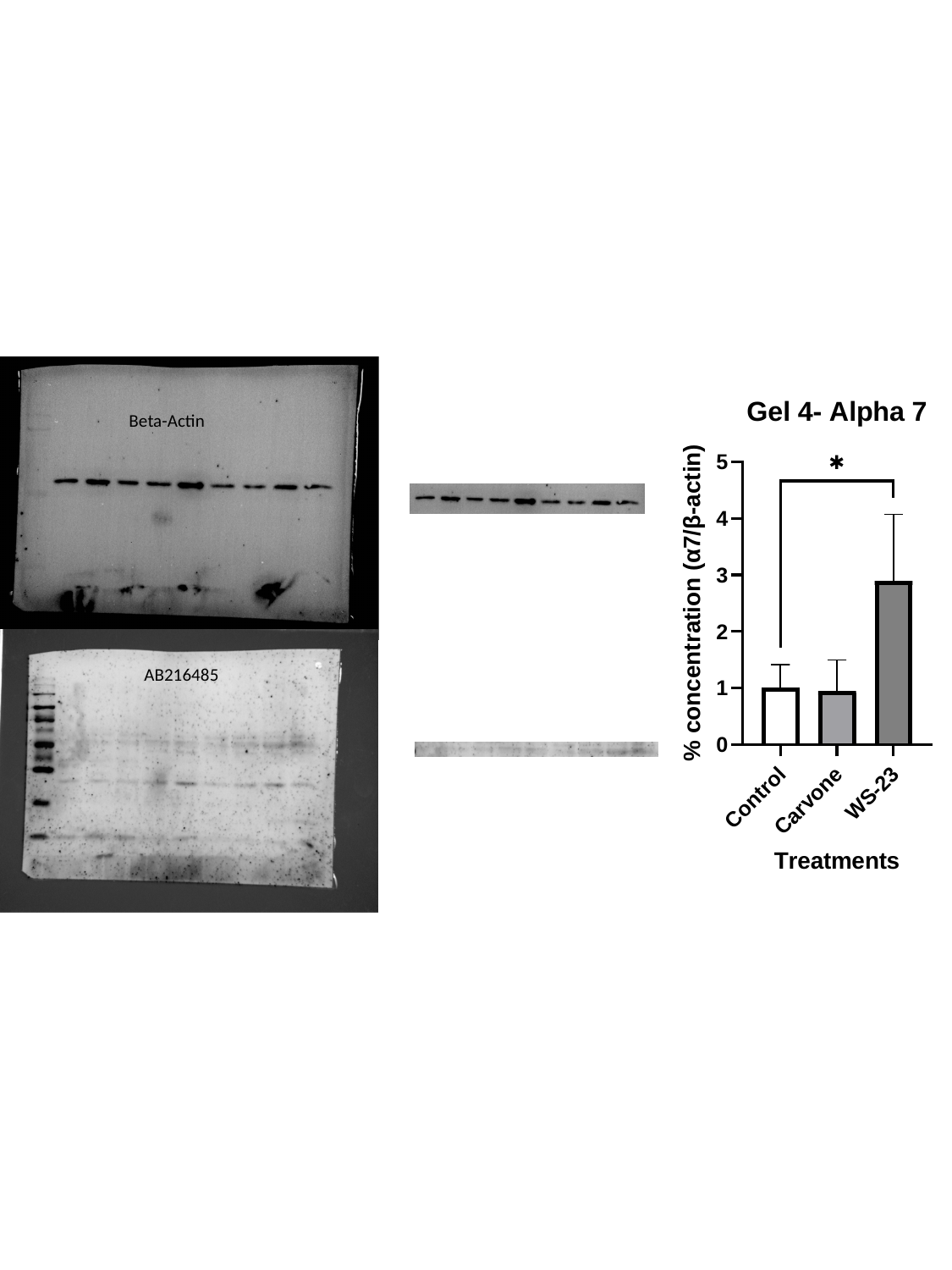

Beta-Actin
AB216485

### Slide 13
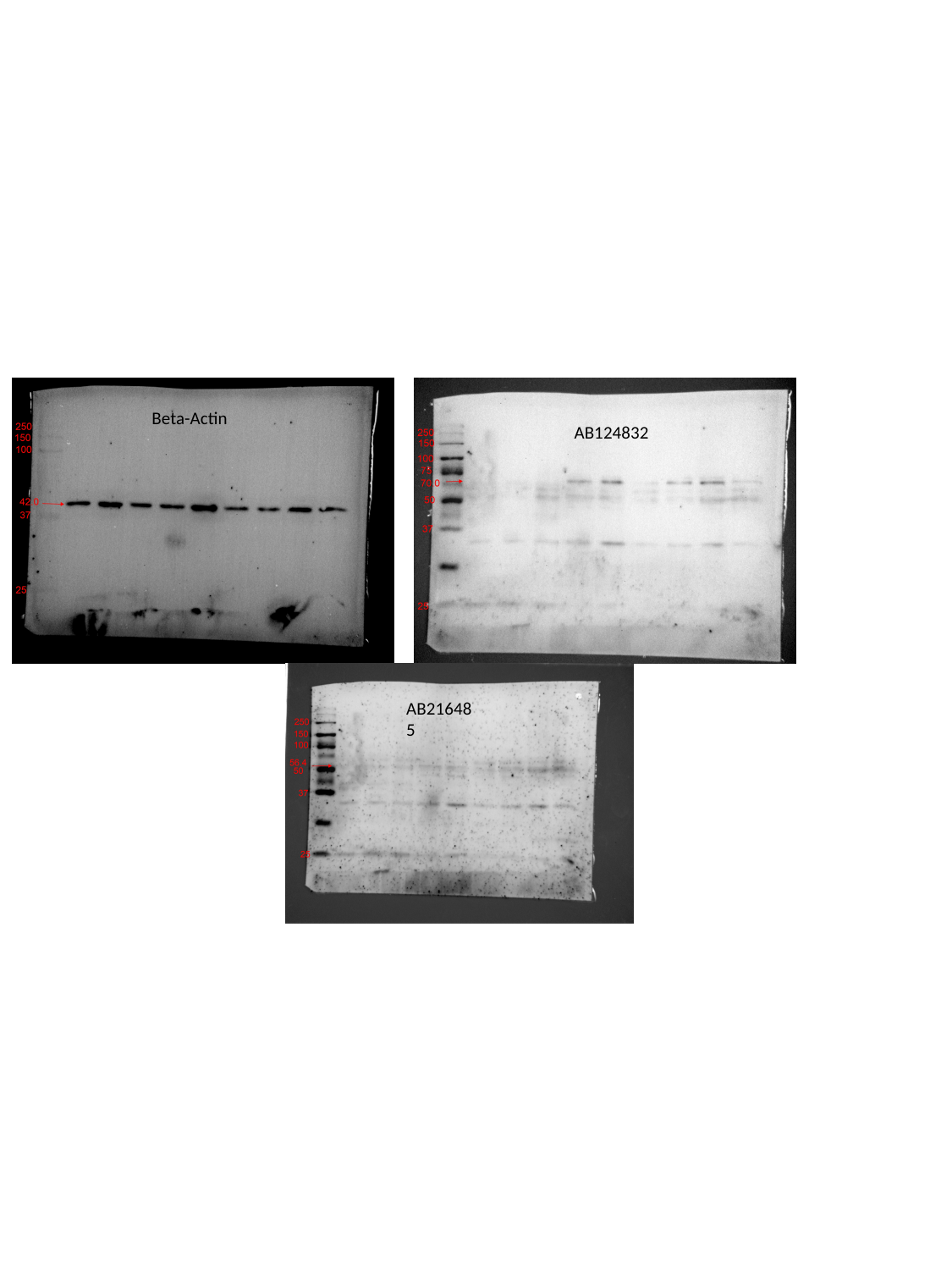

Beta-Actin
AB124832
AB216485
